# Mechanisms of γ-Secretase Activation and Substrate Processing

**DOI:** 10.1101/2020.03.09.984534

**Authors:** Apurba Bhattarai, Sujan Devkota, Sanjay Bhattarai, Michael S. Wolfe, Yinglong Miao

## Abstract

Amyloid β-peptide, the principal component of characteristic cerebral plaques of Alzheimer’s disease (AD), is produced through intramembrane proteolysis of the amyloid precursor protein (APP) by γ-secretase. Despite the importance in pathogenesis of AD, the mechanisms of intramembrane proteolysis and substrate processing by γ-secretase remain poorly understood. Here, complementary all-atom simulations using a robust Gaussian accelerated molecular dynamics (GaMD) method and biochemical experiments were combined to investigate substrate processing of wildtype and mutant APP by γ-secretase. The GaMD simulations captured spontaneous activation of γ-secretase, with hydrogen bonded catalytic aspartates and water poised for proteolysis of APP at the ε cleavage site. Furthermore, GaMD simulations revealed that familial AD mutations I45F and T48P enhanced the initial ε cleavage between residues Leu49-Val50, while M51F mutation shifted the ε cleavage site to the amide bond between Thr48-Leu49. Detailed analysis of the GaMD simulations allowed us to identify distinct low-energy conformational states of γ-secretase, different secondary structures of the wildtype and mutant APP substrate, and important active-site sub-pockets for catalytic function of the enzyme. The simulation findings were highly consistent with experimental analyses of APP proteolytic products using mass spectrometry and western blotting. Taken together, the GaMD simulations and biochemical experiments have enabled us to elucidate the mechanisms of γ-secretase activation and substrate processing, which should facilitate rational computer-aided drug design targeting this functionally important enzyme.

## Introduction

Alzheimer’s disease (AD) is a neurodegenerative disorder characterized by cerebral atrophy, beginning with areas of the brain involved in learning and memory. Deposition of 42-residue amyloid β-peptide (Aβ42) in the form of plaques is a defining pathological feature of AD and begins many years before onset of symptoms.^1^ For these reasons, Aβ42 has been a major target for the development of potential therapeutics^2^ as well as a key biomarker for AD.^3^ Aβ peptides are derived through proteolytic processing of the membrane-traversing amyloid precursor protein (APP), first by β-secretase outside the membrane, generating a membrane-bound 99-residue C-terminal fragment (C99), and then by γ-secretase within the membrane.^4^ γ-Secretase is a membrane-embedded aspartyl protease complex, with presenilin (PS1) as the catalytic component that carries out intramembrane proteolysis of >90 substrates, including APP and the Notch family of cell-surface receptors.^5^ Cleavage of the APP transmembrane (TM) domain by γ-secretase determines the length of Aβ peptides, the proportion of the hydrophobic TM domain retained in the Aβ product, and therefore the tendency of Aβ to aggregate into plaques.

Proteolysis of the APP TM domain by γ-secretase is complex.^6^ Initial endoproteolysis of C99 at the ε site generates 48- or 49-residue Aβ (Aβ48 or Aβ49) and corresponding APP intracellular domains (AICD49-99 or AICD50-99) **(Figure S1)**.^7^ These initially formed Aβ peptides are then trimmed every 3-4 amino acids through a carboxypeptidase activity of γ-secretase along two pathways, Aβ48→Aβ45→Aβ42→Aβ38 and Aβ49→Aβ46→Aβ43→Aβ40,^8-9^ and this trimming is dictated by three active-site pockets that recognize substrate residues P1’, P2’ and P3’ (i.e., immediately C-terminal of the scissile amide bond).^10^ Mutations in the APP TM domain associated with early-onset familial AD (FAD) can skew ε cleavage in favor of Aβ48 (i.e., to the pathological Aβ42 pathway).^10-11^ Alternatively, these mutations can be “pathway switchers”, affecting carboxypeptidase activity to switch from the Aβ40 pathway to the Aβ42 pathway.^10^

Little is known about the mechanism by which γ-secretase accomplishes intramembrane proteolysis. A substantial advance in understanding substrate recognition came recently with reports of cryo-electron microscopic (cryo-EM) structure determination of the γ-secretase complex bound to the Notch and APP substrates **(Figure S1)**.^12-13^ The cryo-EM structures were consistent with expectations from previous studies using small-molecule probes and mutagenesis. In both structures, the substrate TM assumed a helical conformation starting from the extracellular side and was surrounded by TM2, TM3 and TM5 of PS1. The helix ended just before entry into the enzyme active site, becoming first partially unwound and then fully extended into a β-strand toward the intracellular side. The substrate β-strand interacted with an antiparallel β-strand in the intracellular side of PS1 TM7, which in turn interacted with another β-strand from the enzyme TM6. This β-sheet motif was suggested to be essential for substrate recognition by the γ-secretase.^13-14^ While a *tour de force* for the field, stabilization of the substrate-enzyme complex required (1) mutation of one of the catalytic aspartates (Asp385) to alanine in PS1 (inactivating the enzyme) and (2) double cysteine mutagenesis and disulfide crosslinking between substrate and presenilin (with the potential for deviation from normal wildtype interactions).

Computational modeling, especially molecular dynamics (MD) simulation, has proven useful in understanding the structural dynamics of γ-secretase. Previous studies have provided valuable insights into the conformational changes^15-18^, enzyme allosteric modulation^19^, substrate binding^15, 18, 20-22^, water distribution^15-16^, lipid interactions^16^ and ligand binding of γ-secretase^23-25^. A putative active conformation was described for the substrate-free (apo) γ-secretase with the two catalytic aspartates moving to close proximity in few^15-17^, but none has characterized the enzyme active state poised for proteolysis with both the water and peptide substrate. Hence, the dynamic mechanisms of enzyme activation and substrate processing by γ-secretase remained poorly understood.

Here, we present the first report of MD computational modeling of activation of APP-bound γ-secretase using the latest cryo-EM structures of substrate-bound enzyme. The enzyme and substrate were computationally restored to the wildtype. Extensive all-atom simulations using a novel and robust Gaussian accelerated molecular dynamics (GaMD) method were employed to capture the extremely slow motions underlying activation of γ-secretase for proteolysis of substrate within the cell membrane (k_cat_ in proteoliposomes estimated at 1.9 h^-1^).^26^

GaMD is an enhanced sampling computational technique that works by adding a harmonic boost potential to smooth the biomolecular potential energy surface.^27^ GaMD greatly reduces energy barriers and accelerates biomolecular simulations by orders of magnitude.^28^ GaMD does not require pre-defined collective variables or reaction coordinates. Compared with the enhanced sampling methods that rely on careful selection of the collective variables, GaMD is of particular advantage for studying complex biological processes^29^ such as enzyme activation and substrate processing by γ-secretase. Moreover, because the boost potential follows a Gaussian distribution, biomolecular free energy profiles can be properly recovered through cumulant expansion to the second order.^27^ GaMD builds on the previous accelerated MD (aMD) method^30-31^, but solves its energetic reweighting problem^32^ for free energy calculations of large biomolecules. GaMD has successfully revealed physical pathways and mechanisms of protein folding and ligand binding, which are consistent with experiments and long-timescale conventional MD simulations.^27, 33-34^ It has also been applied to characterize protein-protein,^35-36^ protein-membrane,^37^ and protein-nucleic acid^38-39^ interactions. Therefore, GaMD was applied in this study for enhanced sampling of the γ-secretase complex, a well-known slow enzyme.^40-41^

Furthermore, the GaMD simulations were highly consistent with parallel mass spectrometry (MS) and western blotting biochemical experiments on the processing of both wildtype and mutant APP substrates. Remarkably, one of the mutations (M51F) in APP shifted the substrate ε cleavage site to the amide bond between residue Thr48-Leu49, while another two mutations (I45F and T48P) enhanced the ε cleavage between Leu49-Val50 compared with the wildtype. The GaMD simulations and biochemical experiments together offered a deep atomic-level understanding of intramembrane proteolysis by γ-secretase.

## Results

### Activation of computationally restored wildtype γ-secretase is captured in GaMD simulations

Our initial testing GaMD simulations using the earlier published cryo-EM structure of Notch-bound γ-secretase **(Figure S1A)—**with Asp385 computationally restored**—** showed that Asp257 rather than Asp385 should be protonated in the active site, as in this case the two aspartates were able to approach each other for catalysis **(Table 1** and **Figure S2)**. Further testing GaMD simulations using the cryo-EM structure of APP-bound γ-secretase **(Figure S1B** and **Table 1)** revealed an active conformation of the PS1 catalytic subunit with computationally restored Asp385 while the enzyme-substrate disulfide bond was kept **(Figure S3** and **Movie S1)**. Building upon these testing results, we proceeded to remove the artificial enzyme-substrate disulfide bond to completely restore the wildtype γ-secretase for further simulations **(Table 1)**. During three 2-µs GaMD enhanced simulations, spontaneous activation of APP-bound γ-secretase was observed starting from its inactive cryo-EM conformation **(Figures 1A** and **S4A** and **Movie S2)**. The activation was characterized by coordinated hydrogen bonding interactions between the active-site aspartates, APP and a water molecule. Active site Asp257 and Asp385 moved closer to form a hydrogen bond between the protonated Asp257 and the carbonyl oxygen in Leu49 of the scissile amide bond in APP **(Figure 1B)**. The two aspartates were ∼7 Å apart between their Cγ atoms. Water entered the enzyme active site from the intracellular side and formed hydrogen bonds with the aspartates. The hydrogen bonds with the catalytic aspartates activated the water needed for nucleophilic attack of the carbonyl carbon of the scissile amide bond in APP. The distance between the carbonyl carbon of Leu49 and water oxygen was ∼3.8 Å. This active conformation is well poised for ε cleavage of the amide bond between residues Leu49 and Val50 of APP.

**Table 1:** Summary of GaMD simulations performed on different systems of γ-secretase bound by the Notch and APP substrates.

| Enzyme | Substrate | <sup>a</sup>Disulfide Bond | <sup>b</sup>$N_{\text{atoms}}$ | Dimension ($\text{\AA}^3$) | Simulation (ns) | <sup>c</sup>$\Delta V_{\text{avg}}$ (kcal/mol) | <sup>d</sup>$\sigma_{\Delta V}$ (kcal/mol) |
| --- | --- | --- | --- | --- | --- | --- | --- |
| **D385A (Cryo-EM)** | Notch | Present | 240,021 | 141 x 124 x 146 | 300 x 1 | 12.91 | 7.91 |
| **D385-protonated** | Notch | Present | 240,358 | 141 x 124 x 146 | 300 x 3 | 9.97 | 6.76 |
| **D257-protonated** | Notch | Present | 240,358 | 141 x 124 x 146 | 300 x 3 | 10.36 | 6.46 |
| **D385A (Cryo-EM)** | APP | Present | 253,650 | 141 x 124 x 147 | 300 x 1 | 12.53 | 6.58 |
| **Wildtype** | APP | Present | 253,647 | 141 x 124 x 147 | 300 x 3 | 10.46 | 6.91s |
| **Wildtype** | APP | Absent | 241,351 | 141 x 124 x 147 | 2000 x 3 | 10.45 | 6.78 |
| **Wildtype** | I45F APP | Absent | 241,355 | 141 x 124 x 147 | 1100 x 3 | 10.30 | 6.79 |
| **Wildtype** | T48P APP | Absent | 241,348 | 141 x 124 x 147 | 1400 x 3 | 10.87 | 6.87 |
| **Wildtype** | M51F APP | Absent | 241,360 | 141 x 124 x 147 | 1500 x 3 | 10.08 | 7.38 |

<sup>a</sup>The artificial disulfide bond between the N-terminus of APP and PS1 HL1 loop of the $\gamma$ -secretase is kept (“Present”) or removed (“Absent”).
<sup>b</sup> $N_{\text{atoms}}$ is the number of atoms in the simulation systems.
<sup>c</sup> $\Delta V_{\text{avg}}$ and <sup>d</sup> $\sigma_{\Delta V}$ are the average and standard deviation of the GaMD boost potential, respectively.

**Figure 1:**
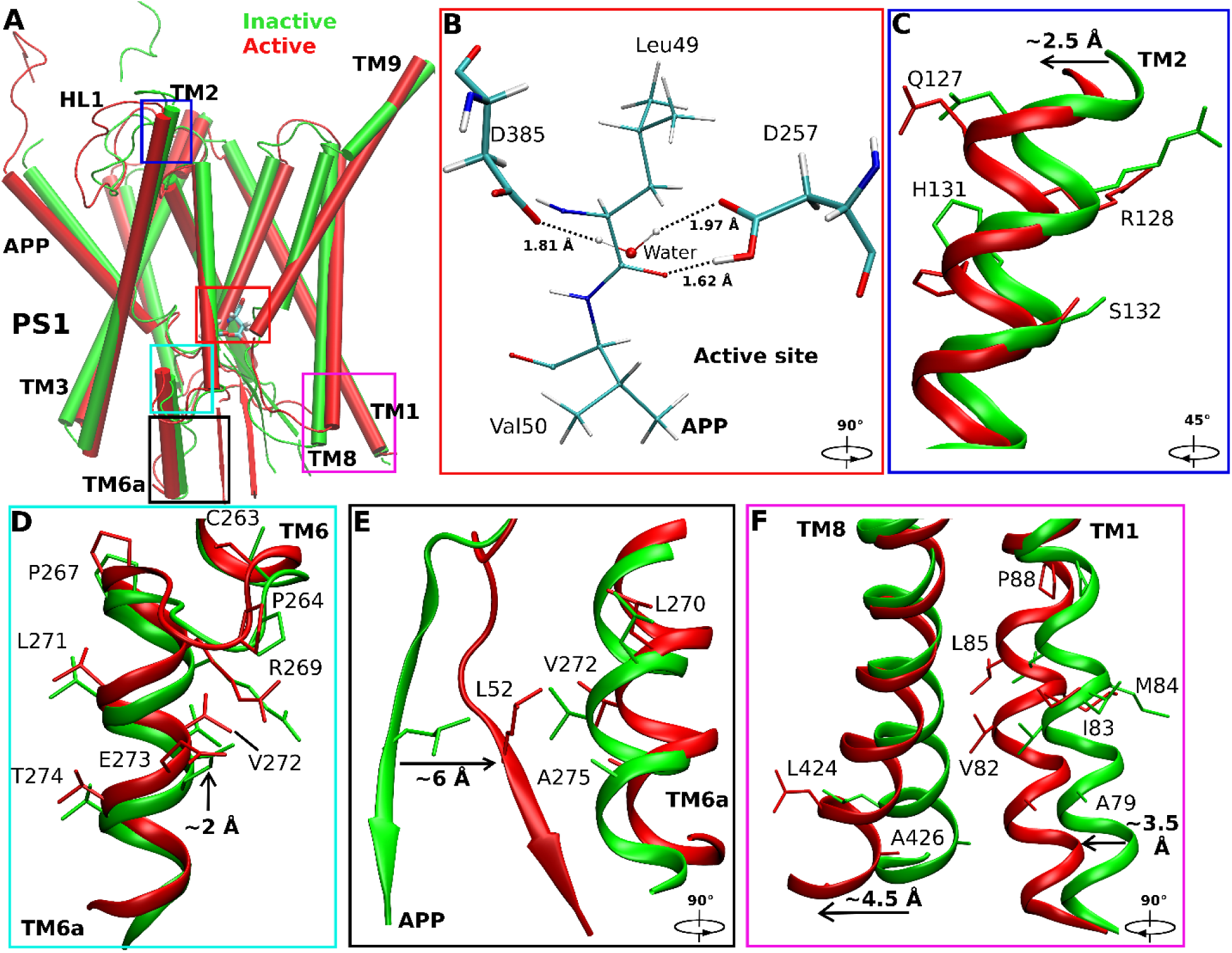
Conformational changes of the catalytic subunit presenilin (PS1) and APP substrate during activation of the computationally restored wildtype γ-secretase. (A) Comparison of the inactive cryo-EM structure (green) and wildtype active conformation of APP-bound PS1 (red). (B) The active site poised for proteolysis. Water entered the active site and formed hydrogen bonds with the catalytic aspartates, being ready for nucleophilic attack on the scissile amide bond between residues Leu49 and Val50 of APP for ε cleavage. (C-F) Conformational changes of (C) PS1 TM2, (D) PS1 TM6a, (E) the C-terminus of APP, (F) PS1 TM1 and PS1 TM8 during activation of γ-secretase. The extracellular end of TM2 moved outwards by ∼2.5 Å in the active PS1 relative to the inactive cryo-EM structure. The PS1 TM6a moved upwards by ∼2 Å compared to the cryo-EM structure. The C-terminal β-strand region of APP moved closer to interact with the PS1 TM6a helix. Residue Leu52 of APP moved by ∼6 Å towards non-polar residues Val272, Leu270 and Ala275 in the enzyme TM6a. The intracellular ends of TM8 and TM1 moved from the cryo-EM structure by ∼4.5 Å and ∼3.5 Å, respectively.

RMSFs were calculated from GaMD simulations of the enzyme-substrate complex **(Figure S5)**. In nicastrin, extracellular helices α1, α2, α4a, the C-terminal regions of α5, α12, α17, and TM domain exhibited high fluctuations with ∼3 Å RMSF **(Figure S5A)**. The TM6 and Helix-8 of APH1 were also flexible during the simulations. In PS1, TM2 extracellular domain, TM6 and TM6a were flexible with ∼2.5-3 Å RMSF. Through structural clustering of GaMD simulation snapshots **(see Methods)**, the top cluster was obtained as the representative wildtype active conformation of the enzyme. Relative to the cryo-EM structure, the extracellular end of TM2 moved outwards by ∼2.5 Å **(Figure 1C)** and TM6a moved upwards by ∼2 Å **(Figure 1D)**. Conformational changes of these domains involved a significant number of PS1 FAD mutation sites, including Gln127, Arg128, Ser132, Pro264, Pro267, Arg269, Leu271, Val272, Glu273 and Thr274 (www.alzforum.org). Interestingly, His131 from TM2 and Cys263 from TM6a flipped their side chains. The N-terminal helix region of APP moved outwards by ∼10 Å during enzyme activation (**Figure 1A**), while the C-terminal β-strand of APP moved by ∼6 Å to interact with the PS1 TM6a helix. In the process, APP residue Leu52 made new contacts with residues Val272 and Ala275 in TM6a of PS1 **(Figure 1E)**. The movement was consistent with previous finding that TM6a undergoes large conformational change upon substrate binding and plays a key role in activation of the enzyme.^12^ In addition, the intracellular ends of TM8 and TM1 moved by ∼4.5 Å and ∼3.5 Å, respectively (**Figure 1F**). Residues Ser104, Phe105 and Tyr106 in the N-terminal region of PS1 HL1 changed into a helical conformation during enzyme activation **(Figures S6B** and **S6F)**. In summary, we have captured activation of computationally restored wildtype γ-secretase bound by wildtype APP in the GaMD simulations.

### GaMD simulations correlated with biochemical experiments on cleavage of wildtype and mutant APP

MS experiments were carried out to analyze AICD species (AICD49-99 and AICD50-99) generated in proteolysis of the wildtype APP and three mutants (I45F, T48P and M51F) by γ-secretase assay **(Figures 2A-2D)**. For the wildtype APP, the MALDI-TOF analysis showed the presence of both AICD species, but the AICD50-99 species had relatively higher intensity than the AICD49-99 species **(Figure 2A)**. The difference in the amount of AICD fragments suggested that the γ-secretase preferred ε cleavage between Leu49-Val50 to the cleavage between Thr48-Leu49 in the wildtype APP, as has been previously reported.^10^ Such experimental data correlated well with GaMD simulations of γ-secretase with the wildtype APP substrate, during which the activated enzyme was poised to cleave wildtype APP between Leu49-Val50 **(Figure 1B)**.

**Figure 2:**
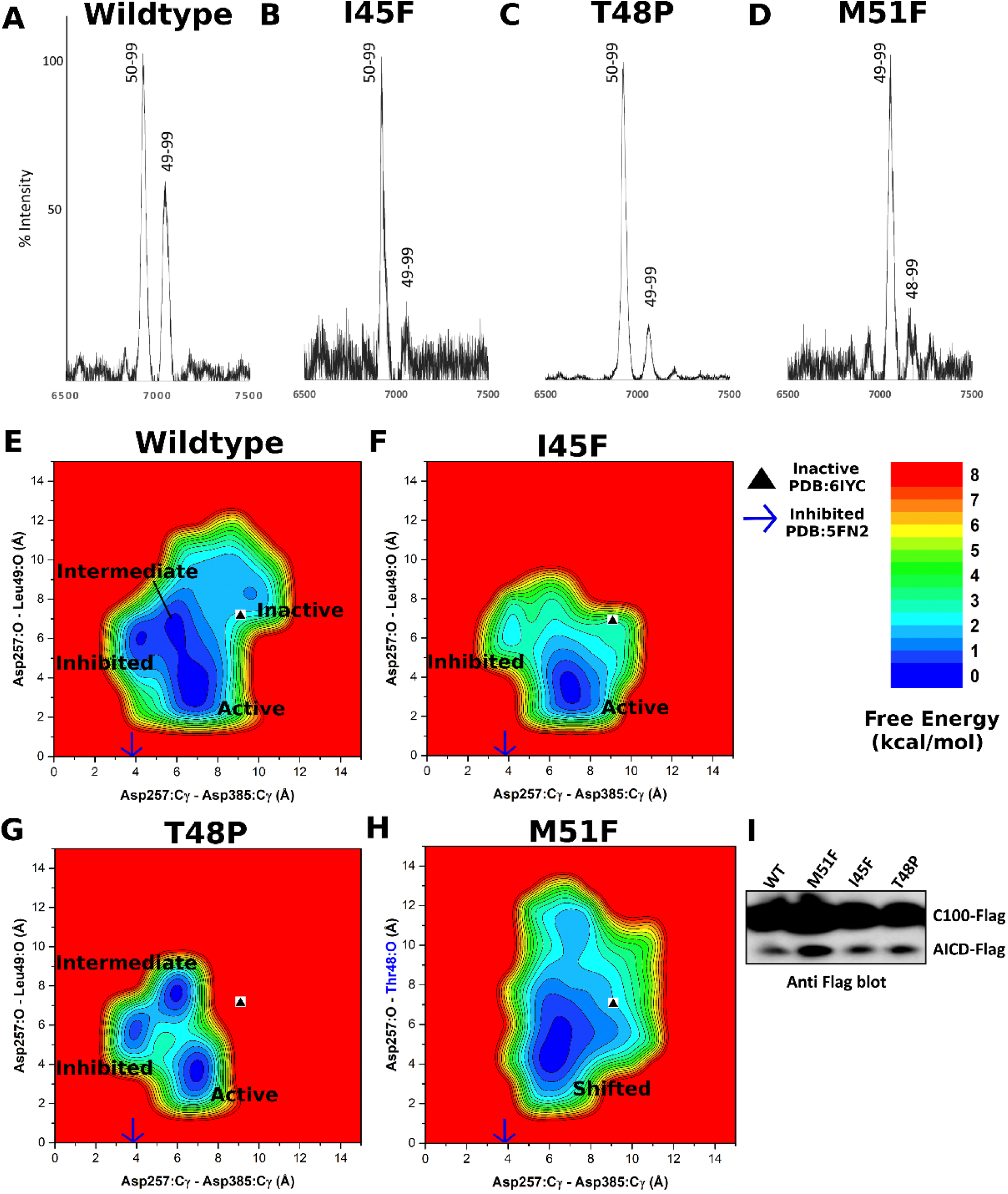
Mass spectrometry and western blotting of the APP intracellular domain (AICD) fragments and GaMD free energy profiles of wildtype and mutant APP-bound γ-secretase. (A-D) The intensity of different AICD fragments detected by mass spectrometry for (A) wildtype (AICD 50-99, expected mass: 6905.6 g/mol, observed mass: 6907.4 g/mol; AICD 49-99, expected mass: 7018.8 g/mol, observed mass: 7019.6 g/mol), (B) I45F (AICD 50-99, expected mass: 6905.6 g/mol, observed mass: 6905.4 g/mol; AICD 49-99, expected mass: 7018.8 g/mol, observed mass: 7019.8 g/mol), (C) T48P (AICD 50-99, expected mass: 6905.6 g/mol, observed mass: 6907.4 g/mol; AICD 49-99, expected mass: 7018.8 g/mol, observed mass: 7041.8 g/mol) and (D) M51F (AICD 49-99, expected mass: 7034.8 g/mol, observed mass: 7031.4 g/mol; AICD 48-99, expected mass: 7135.8 g/mol, observed mass: 7132.2 g/mol) APP substrate as cleaved by γ-secretase. (E-G) 2D free energy profiles of the Asp257:Cγ - Asp385:Cγ and Asp257:protonated O - Leu49:O distances calculated from GaMD simulations of (E) wildtype, (F) I45F and (G) T48P APP substrate. (H) 2D free energy profile of the Asp257:Cγ - Asp385:Cγ and Asp257:protonated O – Thr48:O distances calculated from GaMD simulations of the M51F APP substrate. (I) The total amount of AICD species in γ-secretase determined *in vitro* by western blotting using anti-Flag antibodies of γ-secretase.

During activation, the wildtype APP-bound γ-secretase also sampled “inhibited’ and “intermediate” low-energy states as identified from the GaMD reweighted free energy profile **(Figure 2E)**. In the inhibited state, the catalytic aspartates moved very close to each other, with only ∼4 Å distance between the Cγ atoms, while the substrate was ∼6 Å away from the active site. This conformation could not accommodate water between the aspartates to form hydrogen bonds. A similar inhibited state of the enzyme was also observed in the dipeptidic inhibitor N-[N-(3,5-difluorophenacetyl)-L-alanyl]-S-phenylglycine t-butyl ester (DAPT)-bound cryo-EM structure (PDB: 5FN2).^42^ With the enzyme active site in the inhibited state, APP substrate moved away from the catalytic aspartates in the GaMD simulations. The carbonyl oxygen in Leu49 of APP was ∼6 Å from the protonated oxygen of Asp257 **(Figure 2E)**. In the intermediate state, the Cγ atoms of the catalytic aspartates were ∼6 Å apart, while the carbonyl oxygen of Leu49 of the APP substrate was ∼7 Å away from the Asp257 oxygen **(Figure 2E)**.

For the I45F and T48P mutants of APP substrate, the MS analysis showed decreased amount of the AICD49-99 species from proteolysis of both mutants compared with the wildtype substrate, and AICD50-99 was the predominant AICD product **(Figures 2B and 2C)**. Thus ε cleavage between Leu49-Val50 was even more preferred for these two mutants than for the wildtype substrate. In parallel with the experiments, further GaMD simulations were performed on γ-secretase bound by the I45F and T48P mutant APP substrates. The I45F mutant substrate-bound γ-secretase became activated during 1.1 µs GaMD simulations **(Figure S4B)**. A low-energy conformation was observed in the I45F active state for which the distance between the Cγ atoms of Asp257 and Asp385 was ∼7 Å, while the Leu49 carbonyl oxygen and Asp257 protonated oxygen formed a hydrogen bond with ∼3 Å distance **(Figure 2F)**. The boost potential was 10.30 ± 6.79 kcal/mol in the GaMD simulations of the I45F mutant substrate-bound enzyme, which was comparable to that of the wildtype system (10.45 ± 6.78 kcal/mol) **(Table 1)**. However, the I45F-mutant APP substrate-bound γ-secretase was activated within shorter simulation time compared with the wildtype APP substrate-bound enzyme, with higher probability of conformations for the ε cleavage between Leu49-Val50 in APP **(Figures S4A and S4B)**. The simulation findings agreed well with the experimental data. Analysis of AICD products by MALDI mass spectrometry revealed a higher peak intensity of AICD50-99 than AICD49-99 for the I45F mutant APP compared with the wildtype APP **(Figures 2A and 2B)**. Another low-energy conformation of I45F APP substrate-bound enzyme was observed in the inhibited state **(Figure 2F)**, being similar to the inhibitor DAPT-bound structure of γ-secretase (PDB: 5FN2)^42^.

For the T48P-mutant APP substrate-bound γ-secretase, activation was observed during one of three 1.5 µs GaMD simulations **(Figure S4C)**. Low-energy conformations were identified from the free energy profile in the active, inhibited and intermediate states **(Figure 2G)**. In the T48P active state, the catalytic aspartates were positioned ∼7 Å apart (Cγ to Cγ), and the substrate Leu49 carbonyl oxygen aligned with the protonated oxygen of Asp257 to form a hydrogen bond. The boost potential was 10.87 ± 6.87 kcal/mol, which was also comparable to that of wildtype APP simulations **(Table 1)**. The T48P-mutant APP substrate-bound γ-secretase transitioned into the active state within shorter simulation time compared to the wildtype system **(Figure S4A and S4C)**. The T48P APP substrate mutant had a higher probability than the wildtype APP substrate of aligning the aspartates and water with the scissile amide bond between Leu49-Val50 in APP. This computational finding was again consistent with MALDI mass spectrometric analysis of AICD products: AICD50-99 intensity is higher than AICD49-99 for the T48P-mutant substrate compared to that of wildtype system **(Figures 2A and 2C)**. The observed inhibited state (**Figure 2G**) was similar to that seen in the wildtype and I45F systems **(Figures 2E and 2F)** as well as the inhibitor DAPT-bound cryo-EM structure of γ-secretase (PDB: 5FN2).^42^ In the T48P intermediate state, the Cγ atoms of the catalytic aspartates were ∼6 Å apart, while the Leu49 carbonyl oxygen of APP substrate and the protonated oxygen of Asp257 were ∼7 Å apart **(Figure 2G)**. The I45F and T48P-mutant APP substrate-bound γ-secretase showed similar structural flexibility as the wildtype system in the RMSFs calculated from GaMD simulations **(Figures S5A)**. In both systems, extracellular helices α1, α2, α4a, the C-terminal regions of α5, α12, α17, and TM domain of nicastrin exhibited high fluctuations. The TM6 and Helix-8 of APH-1, and the TM2 extracellular domain, TM6 and TM6a of PS1 were also flexible during the GaMD simulations **(Figures S5C and S5D)**.

### Shifted ε cleavage site of APP in the M51F mutant

MS analysis of AICD products from the M51F-mutant system revealed AICD49-99 as the major product, suggesting that the predominant ε cleavage site of M51F APP was between residues Thr48-Leu49 **(Figure 2D)**. A low level of AICD48-99 was also detected, revealing that the M51F APP substrate was cleaved to a limited degree between Ile47-Thr48 **(Figure 2D)**. This was consistent with previous studies that a Phe residue is not tolerated in the P2’ position of substrate or transition-state analogue inhibitors of γ-secretase.^10, 43^ Thus, M51F mutation of the APP substrate shifted the ε cleavage site from Leu49-Val50 to Thr48-Leu49. Such shift of ε cleavage was consistently observed in 1.5 µs GaMD simulations of the M51F-mutant APP substrate bound to γ-secretase **(Figure S4D** and **Movie S3)**. The protonated oxygen of PS1 Asp257 was hydrogen bonded to the carbonyl oxygen of Thr48, and the activated water molecule targeted the scissile amide bond between Thr48 and Leu49 in the M51F APP mutant for ε cleavage. In comparison, residue Thr48 in the wildtype APP maintained a distance of ∼8-9 Å between its carbonyl oxygen and the protonated oxygen of the PS1 Asp257 **(Figure S4E)**. A distinct low-energy state was identified for the “shifted” conformation in the free energy profile of the M51F APP system **(Figure 2H)**. In the shifted state, the Cγ atoms of the catalytic aspartates were ∼7 Å apart, and the carbonyl oxygen of APP substrate Thr48 and protonated oxygen of PS1 Asp257 formed a hydrogen bond with ∼3 Å distance. Moreover, in one of the three GaMD simulations, the ε cleavage site of M51F-mutant APP was further shifted to the amide bond between Ile47-Thr48. The distance between the carbonyl oxygen of APP substrate residue Ile47 and the protonated oxygen of PS1 Asp257 became ∼3 Å **(Figure S7)**. The Cγ atoms of the catalytic aspartates were ∼7 Å apart. This observation was consistent with the low level of AICD48-99 fragment detected by MS of AICD products of the M51F APP mutant **(Figure 2D)**.

In addition to the MS experiments, the effect of APP mutations was investigated by detecting the total amount of Flag-tagged AICD species in *in-vitro* γ-secretase assays by western blotting using anti-Flag antibodies (**Figure 2I)**. The AICD production increased substantially for the M51F mutant substrate compared to wildtype APP substrate. In contrast, ε proteolysis of the I45F and T48P mutants showed no drastic change in the total AICD level compared with wildtype APP substrate **(Figure 2I)**. This was highly consistent with the GaMD simulations. In the systems with wildtype, I45F and T48P mutant APP substrate, the low-energy inhibited state was observed in the free energy profiles of γ-secretase, but not for M51F-mutant system **(Figures 2E-2H)**.

Structural clustering was performed on GaMD simulations of M51F APP-bound γ-secretase, and the top cluster was identified as the shifted conformational state of the enzyme. Compared with the wildtype active conformation **(Figure 3A)**, the extracellular end of TM2 in PS1 moved outwards by ∼2.5 Å **(Figure 3B)**. The helix involving residues Thr124, Val125, Gly126 and Gln127 became disordered in this region **(Figure 3B)**. The APP substrate moved downwards by ∼4 Å in the substrate binding channel of the enzyme **(Figures 3A and 3C)**. In comparison, the catalytic aspartates and flanking regions of TM6 and TM7 moved less than APP **(Figures 3C and 3E)**. Upon shifting of the ε cleavage site, local rearrangements of APP residues were required to establish the coordinated hydrogen bonding interactions at the active site **(Figure 3C** and **Movie S3)**. Sidechain flipping of the APP Thr48 residue led to formation of a hydrogen bond between its carbonyl oxygen and the PS1 Asp257 protonated oxygen (**Figure S4D**). Residue Leu49 initially facing the activated water between these catalytic aspartates flipped the sidechain and moved downwards by ∼4 Å. The PS1 TM6 helix in the M51F shifted state moved towards the active site by ∼4 Å relative to the wildtype active conformation **(Figure 3A)**. Moreover, the TM6a helix tilted by ∼60° relative to the wildtype active conformation **(Figures 3A and 3D)**. Meanwhile, the β-strand at the C-terminus of APP deformed to a turn as it moved away from the TM6a helix. APP Leu52, interacting with the non-polar residues of TM6a in the wildtype active conformation, flipped its side chain and moved away from these residues by ∼6 Å **(Figure 3E)**. The intracellular domains of TM1 and TM8 in the M51F shifted state moved by ∼2.5 Å compared with the wildtype active conformation **(Fig 3F)**. PS1 FAD mutation sites Ala79, Val82, Ile83, Met84, Leu85, Pro88, Leu424 and Ala426 from TM1 and TM8 showed similar movements of their sidechains. In addition, RMSF of the M51F-mutant APP-bound γ-secretase calculated from GaMD simulations showed higher flexibility in TM2, TM6 and TM6a regions of PS1 **(Figures S5B)**. This extra flexibility is consistent with the ability of the M51F-mutant system to readjust the positioning of the substrate in the active site in shifting the ε cleavage site.

**Figure 3:**
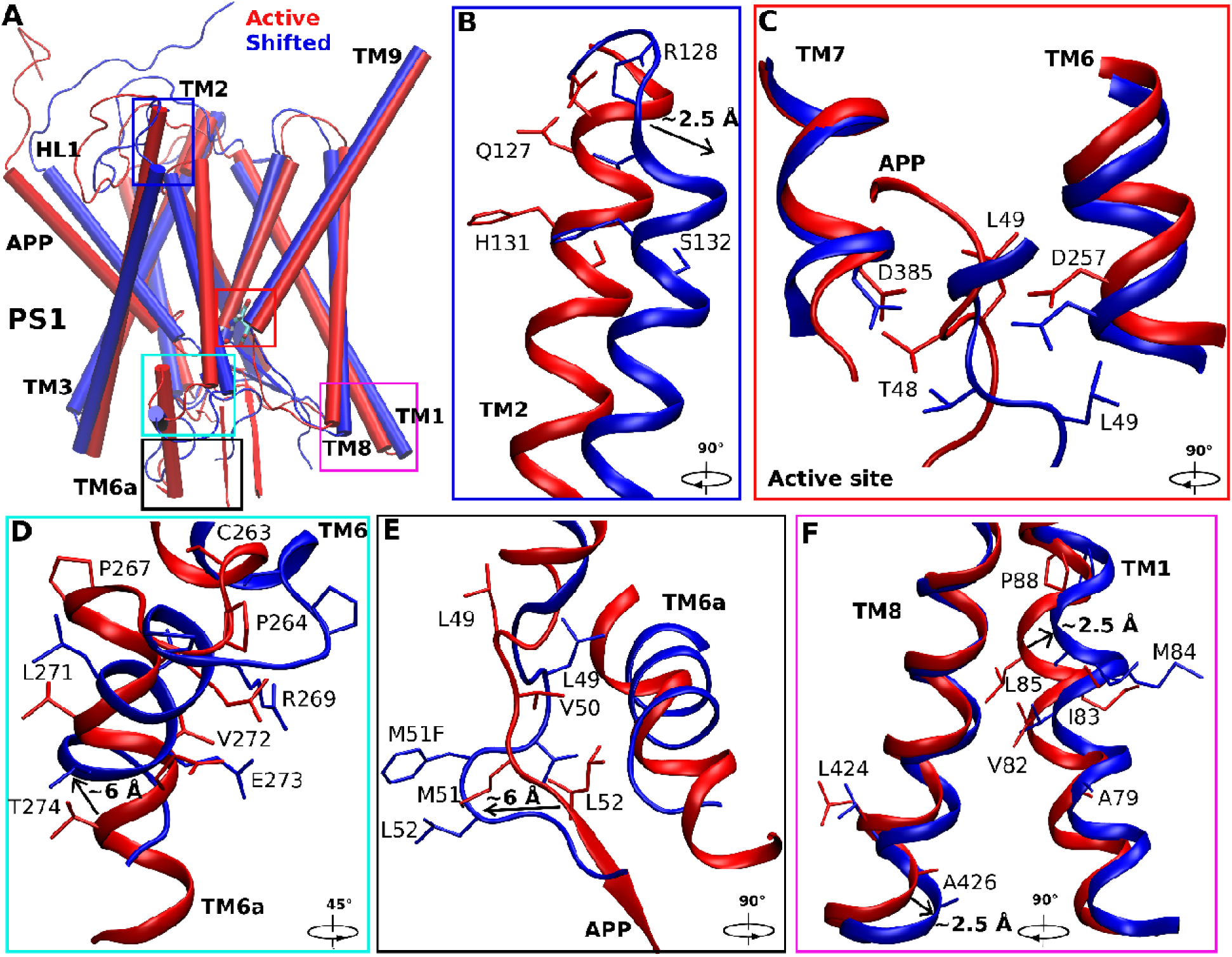
Conformational changes of the catalytic subunit presenilin (PS1) and APP in the shifted active (M51F) states of γ-secretase compared with the active (wildtype) state. (A) Overview of the active (red) and shifted active (blue) conformations of APP-bound PS1. (B) The extracellular end of the PS1 TM2 moved outwards by ∼2.5 Å in the shifted active (M51F) conformation relative to the active (wildtype) structure. Residues Thr124, Val125, Gly126 and Gln127 in this region lost the helical conformation in the shifted active state. (C) The active site poised to attack the scissile amide bond between residues Leu49 and Val50 in the active state (red) and between residues Thr48 and Leu49 in the shifted active state (blue) of APP for ε cleavage. Sidechain flipping of the APP Thr48 residue led to formation of a hydrogen bond between its carbonyl oxygen and the PS1 Asp257 protonated oxygen. Residue Leu49 initially facing the center of two aspartates flipped to the other side with downward movement of ∼4 Å. (D) The N-terminus of PS1 TM6 moved towards the active site by ∼4 Å and the TM6a helix tilted by ∼60°. (E) The β-strand at the C-terminus of APP substrate deformed to a turn as it moved away from TM6a in PS1. The APP Leu52 interacting with non-polar residues in PS1 TM6a in the active conformation flipped its side chain and moved in the opposite direction by ∼6 Å. (F) The intracellular ends of TM1 and TM8 moved by ∼2.5 Å and ∼2.5 Å in PS1, respectively.

### Changes in secondary structures of APP substrate mutants

Changes in secondary structures of the wildtype and mutant APP substrate in γ-secretase were monitored during the GaMD simulations **(Figures 4, S8** and **S9)**. Secondary structures of APP substrate in the active conformations of the wildtype, I45F and T48P mutant systems and the shifted active conformation of the M51F mutant system were compared using their top ranked structural clusters obtained from the corresponding simulations **(Figure S10)**. For wildtype APP substrate, residues Gly29 to Val46 formed a helical conformation throughout the simulations except between residues 42 and 43 **(Figure 4A)**. Residues Asn27-Lys28 fluctuated between turn and coil conformations during last ∼700 ns for activation whereas Ile47-Leu49 fluctuated between helix and turn conformations throughout the simulation. The N-terminal region of APP substrate was very flexible and sampled turn and coil conformations. The C-terminal residues Leu52 to Lys55 primarily maintained an antiparallel β-sheet conformation. Residues Val50-Met51, immediately after the Leu49-Val50 ε cleavage site, formed a turn for a number of times that exposed this APP scissile amide bond to the enzyme aspartates and coordinated water for proteolysis.

**Figure 4:**
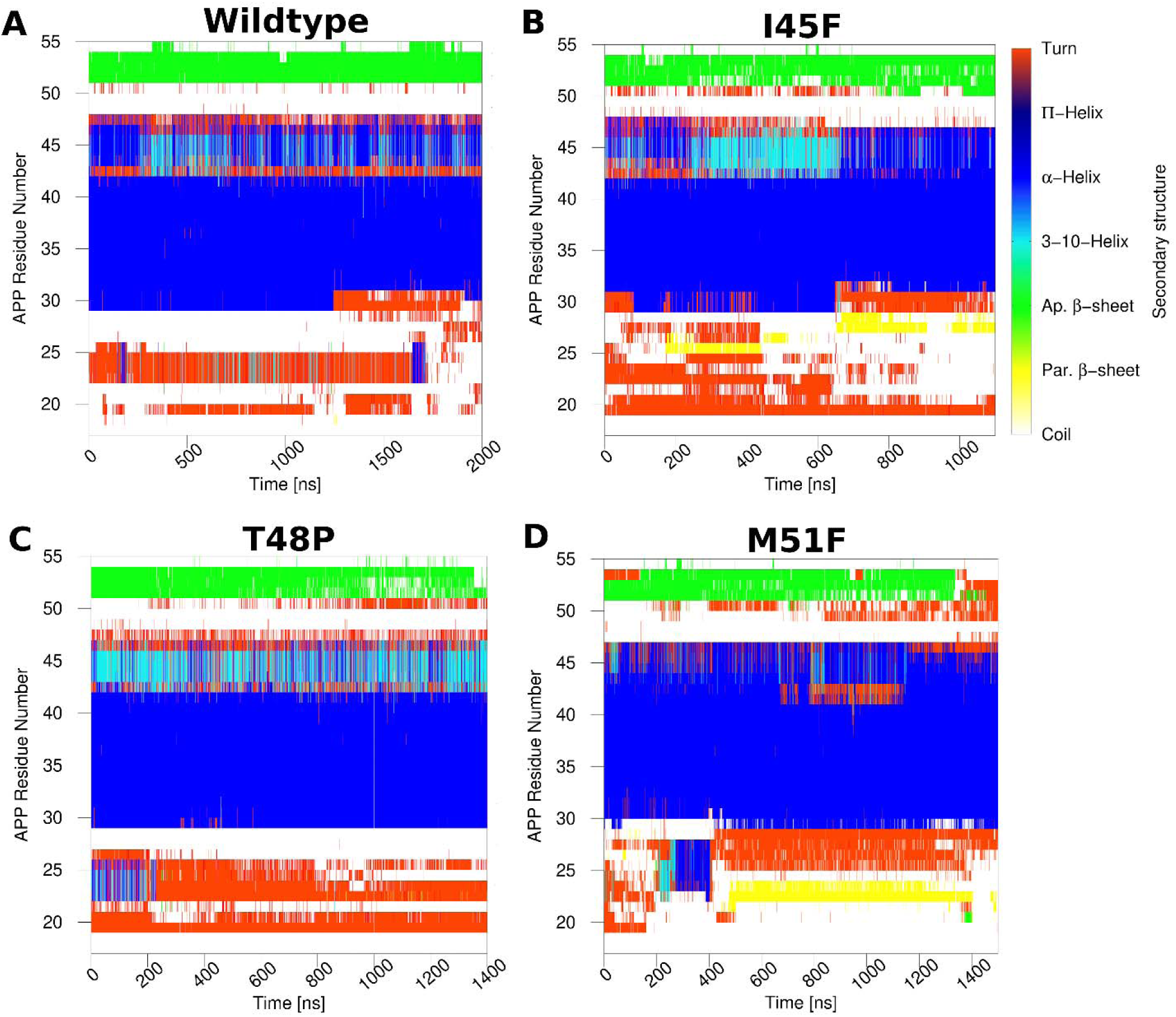
Time courses of the APP secondary structures in the (A) wildtype, (B) I45F, (C) T48P and (D) M51F forms as bound to γ-secretase calculated from their representative GaMD simulations. Results of the other simulations are plotted in **Figures S8** and **S9**.

The I45F and T48P mutants of APP substrate, which maintained ε cleavage between Leu49-Val50, showed similar secondary structures as the wildtype substrate, although unique features were also observed in each mutant. Both mutant substrates formed turns at residues Val50-Met51 despite fluctuations **(Figures 4B** and **4C)** and adopted a β-sheet conformation at the C-terminus during simulations. However, only residues Ile31-Val46 formed a helix in the I45F mutant, with ∼2-3 residues towards the N-terminus losing the helical conformation **(Figure 4B)**. The N-terminal region of I45F-mutant APP substrate was thus more flexible than the wildtype substrate and bent over the HL1 loop of PS1 **(Figures S6C, S6F** and **S11)**. For the I45F APP mutant substrate, multiple hydrogen bonds were formed between the N-terminal residues of APP and PS1 HL1. Residue Gln112 in the PS1 HL1 loop—mutated to Cys112 in the cryo-EM structure to generate a disulfide bond and restored in our simulation—formed three hydrogen bonds with residues Ser26, Asn27 and Lys28 of the APP N-terminus **(Figure S11A)**. Two of these hydrogen bonds involved backbone atoms. In addition, the backbone N atom of Ile114 in PS1 HL1 formed a hydrogen bond with the backbone O atom of Lys28 of APP. These hydrogen bonds contributed to a parallel β-sheet between the PS1 HL1 and APP N-terminus as reflected in the secondary structure plots **(Figures 4B, S8, S11A)**. In contrast, the N-terminal loop of wildtype APP was observed flexible without bending over the PS1 HL1 **(Figures S6B** and **S6F)**. For the T48P-mutant APP, residues Gly29-Ala42 formed a helical conformation, whereas residues Thr43-Ile47 fluctuated between the α-helix, 3-10 helix and turn conformations **(Figure 4C)**. Residues Ser104, Phe105, Tyr106 and Thr107 of the PS1 HL1 loop formed a helix in the T48P active conformation, similar to what was observed in the wildtype active state **(Figure S6D** and **S6F)**.

For the M51F mutant APP, residues Ala30-Val46 formed a helical conformation during the GaMD simulations **(Figure 4D)**. A longer turn appeared starting from residue Leu49 to Leu52 in M51F APP during the simulations **(Figures 4D, 3E** and **S10)**. In comparison, a turn was formed for only residues Val50-Met51 in the wildtype APP that exposed the Leu49-Val50 scissile amide bond for ε cleavage (**Figures 4A** and **S10**). The shift of this turn correlated with the shift of the ε cleavage site. The C-terminal β-strand became shorter in the M51F APP substrate mutant **(Figure 4D)** and even completely disappeared in the representative M51F shifted active conformational state **(Figures S10** and **3E)**. As the M51F-mutant APP substrate moved downwards relative to PS1, its N-terminus formed more interactions with PS1 HL1 **(Figures S6E** and **S6F)**. The backbone O and N atoms of Gly111 in HL1 often formed hydrogen bonds with the backbone atoms N of Glu22 and O of Phe20 in the APP substrate, respectively **(Figure S6E, S6F** and **S11B)**. These hydrogen bonds resulted in a parallel β-sheet conformation **(Figure 4D)**. The N-terminus of the T48P-mutant APP was also found in proximity with the PS1 HL1 loop **(Figure S6D** and **S6F)**. The PS1 HL1 loop—with high flexibility and multiple interactions with APP substrate—make it one of the most important regions of PS1 in the context of enzymatic function and Alzheimer’s disease pathogenesis^44-45^. The PS1 HL1 has a large number of FAD mutation sites, including Phe105, Gly111, Leu113, Tyr115, and Gln127. Hence, our simulation findings were consistent with the literature regarding importance of the HL1 loop.

The C-terminus of APP substrate-bound to the active wildtype conformation moved towards PS1 TM6a region by ∼6 Å during enzyme activation **(Figure 1E)**. The C-terminus of APP maintained β-sheet conformations with the N-terminus of PS1 TM7 throughout the simulations. Hence, with the movement of C-terminus of APP, the N-terminus of PS1 TM7 also moved along by ∼6 Å **(Figure S12)**. In contrast, I45F and T48P mutant APPs maintained the β-sheet conformations with the N-terminus of PS1 TM7 without the movement of C-terminus. M51F mutant APP lost its interaction with the PS1 TM7 and hence losing the β-sheet conformations **(Figures 4D** and **3E)**.

### Comparison of the S1’, S2’ and S3’ active-site subpockets in the wildtype and mutant APP substrate-bound γ-secretase

Representative active conformations of PS1 were identified as the top ranked structural clusters from the GaMD simulations of the wildtype, I45F and T48P mutant systems and the shifted active conformation from the M51F system simulations. These conformations were aligned and compared for the enzyme active-site S1’, S2’ and S3’ subpockets that were occupied by APP substrate residues P1’, P2’ and P3’, respectively^10^ **(Figure 5)**. In the active wildtype conformation, the S1’ subpocket occupied by P1’ residue (Val50) constituted residues mostly from TM6 and TM6a as listed in **Table S1**. The S3’subpocket occupied by P3’ residue (Leu52) constituted residues mostly from TM6a and the C-terminus of PS1-NTF. The S1’ and S3’ subpockets were located on the same side with respect to APP **(Figures 5A)**. In contrast, the S2’ subpocket occupied by P2’ residue (Met51) constituted residues mostly from TM8, TM8-TM9 loop and the β-strand region of TM7 **(Table S3)**.

**Figure 5:**
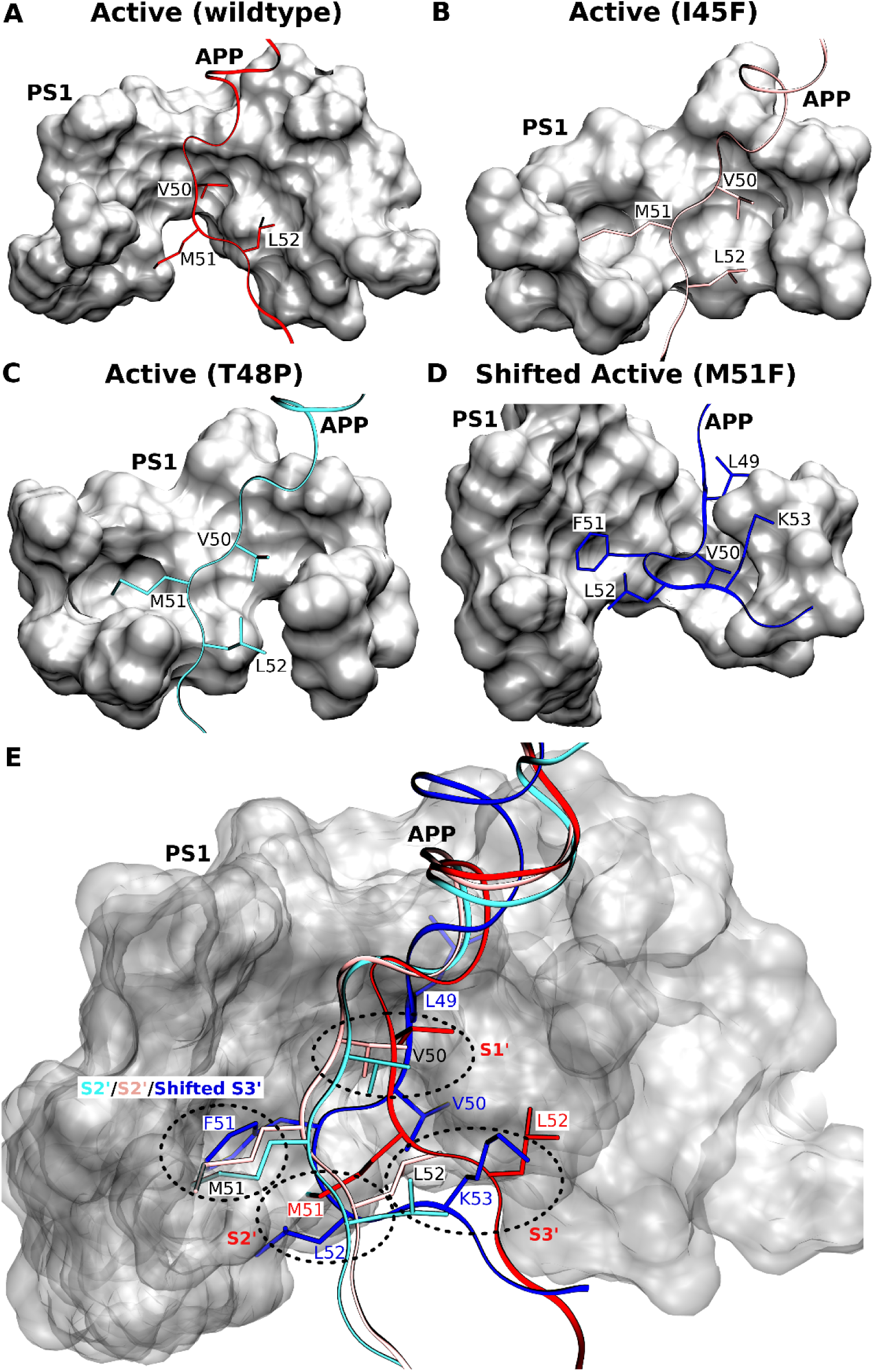
(A-D) Comparison of the locations of APP substrate residues P1’, P2’ and P3’ in the (A) wildtype active, (B) I45F active, (C) T48P active and (D) shifted active M51F APP substrate-bound conformations of γ-secretase. (E) Comparison of the corresponding PS1 active-site S1’, S2’ and S3’ pockets in these different conformational states of γ-secretase.

In the I45F and T48P active conformations **(Figures 5A, 5B and 5C)**, the S1’ and S3’ subpockets occupied by the P1’ (Val50) and P3’ (Leu52) residues, respectively, embodied the same S1’ and S3’ sub pockets of the wildtype active conformation. The S2’ pocket occupied by P2’ (Met51) of the I45F and T48P mutant APP substrate comprised residues from TM8, the TM8-TM9 loop, the β-strand region of TM7 and part of TM1. Notably, both the S1’ and S2’subpockets involved the PAL motif (P333-A434-L435) in the TM9 N-terminal region that is considered important for substrate binding^46^. This S2’ subpocket occupied by the APP mutants was located on the same side of substrate but ∼4 Å above the extended S2’ subpocket in the wildtype active conformation.

For the M51F APP mutant, the presence of a bulky residue Phe at the P2’ position induced local rearrangements and shifted the ε cleavage site. With the shift, Leu49, Val50 and Phe51 became the new P1’, P2’ and P3’ residues, respectively. The new P1’ residue occupied a distinct subpocket near to the S1’ subpocket in the wildtype active conformation **(Figures 5A, 5D** and **5E)**. In contrast, the new P2’ residue occupied a new subpocket in the space between the S1’ and S3’ subpockets of the wildtype active conformation of PS1. The new P3’ residue occupied the same extended pocket as S2’ subpocket in the I45F active and T48P active conformations (**Figures 5B, 5C** and **5E**). Hence, the new subpocket occupied by the P3’ residue (F51) is termed “shifted S3’ subpocket” here and also involved the PAL motif **(Figures 5D** and **5E)**. Moreover, the L52 (P4’) residue and K53 (P5’) residue in the M51F shifted active conformation occupied the S2’ and S3’ subpockets as in the wildtype active conformation of PS1, respectively **(Figures 5D** and **5E)**.

The location of the S2’ subpocket differed among the active conformations of the wildtype active, I45F active and T48P active conformations. As the C-terminus of I45F and T48P mutant APP moved by ∼6 Å compared with the wildtype APP **(Figure S12)**, the P2’ residue (Met51) of these mutants occupied a different S2’ subpocket **(Figures 5)**. Due to shift in the ε cleavage site, the C-terminus of M51F mutant APP lost interactions with the N-terminus of PS1 TM7 and PS1 TM6a. This resulted in large conformational tilting of PS1 TM6a helix in the M51F shifted conformation **(Figure 3)**. Therefore, the conformational changes and molecular interactions of the APP with the γ-secretase provided important insights into the mechanisms of activation and substrate processing by the enzyme.

## Discussion

The PS1-containing γ-secretase complex is a founding member of intramembrane-cleaving proteases (I-CLiPs) which carry out hydrolysis of substrate TM domains within the hydrophobic environment of the lipid bilayer.^47^ I-CLiPs also include the S2P metalloproteases, rhomboid serine proteases, and presenilin-like aspartyl proteases. Although microbial representatives of each of these other I-CLiP classes have been crystallized for high-resolution structure determination ^48-51^, visualizing the active state and elucidating the molecular mechanism of intramembrane proteolysis has been challenging. Only very recently has a rhomboid protease been studied through time-resolved x-ray crystallography to reveal how this serine protease hydrolyzes transmembrane substrates.^52^ Most recently, structures of the γ-secretase complex bound to Notch and APP substrates have been reported, providing critical insights into substrate recognition of γ-secretase.^12, 53^ Nevertheless, mutations in the enzyme and substrate needed for stabilization of the substrate-enzyme complex precluded visualization of the active protease and raised the possibility of unnatural substrate interactions.

Using the latest cryo-EM structures, we have, for the first time, developed an all-atom MD model for activation of the APP substrate-bound γ-secretase poised for intramembrane proteolysis that is in excellent agreement with mass spectrometry and western blotting biochemical experiments. Extensive simulations using a novel GaMD enhanced sampling method have captured spontaneous activation of γ-secretase in the presence of APP and water (**Figure 6**). The catalytic aspartates moved into close proximity, similar to previous simulation findings,^15-17^ although, these studies were performed without the APP substrate bound to the γ-secretase active site. Previous studies suggested a putative active conformation of the apo γ-secretase but was unable to fully characterize the enzyme activation involving additional coordinated hydrogen bond interactions with the substrate. In the GaMD simulations, water molecules entered the active site, one of which coordinated with the two aspartates (**Figure 6B** and **Movies S1** and **S2)**. Moreover, Asp257 formed a hydrogen bond with the carbonyl oxygen of the scissile amide bond between APP residues Leu49-Val50. The activated water molecule was poised for nucleophilic attack on the backbone carbon atom of this activated Leu49-Val50 amide bond. While a number of regions of nicastrin, Aph-1, and Pen-2 displayed flexibility during simulations of the activated enzyme-substrate complex, the PS1 TM6a was the most noteworthy, as this region interacted directly with substrate near the cleavage site and appeared to play a role in enzyme activation. The wildtype enzyme-substrate complex additionally sampled the inhibited and intermediate conformational states, the former closely resembling the conformation of the DAPT inhibitor-bound γ-secretase.^42^ The current ∼2-μs GaMD simulation of γ-secretase with wildtype APP captured the enzyme activation for ε cleavage of APP between Leu49-Val50. The ε cleavage of wildtype APP between Thr48-Leu49 with lower probability, as detected by MS, would likely require longer simulation time and more sufficiently sampling.

**Figure 6:**
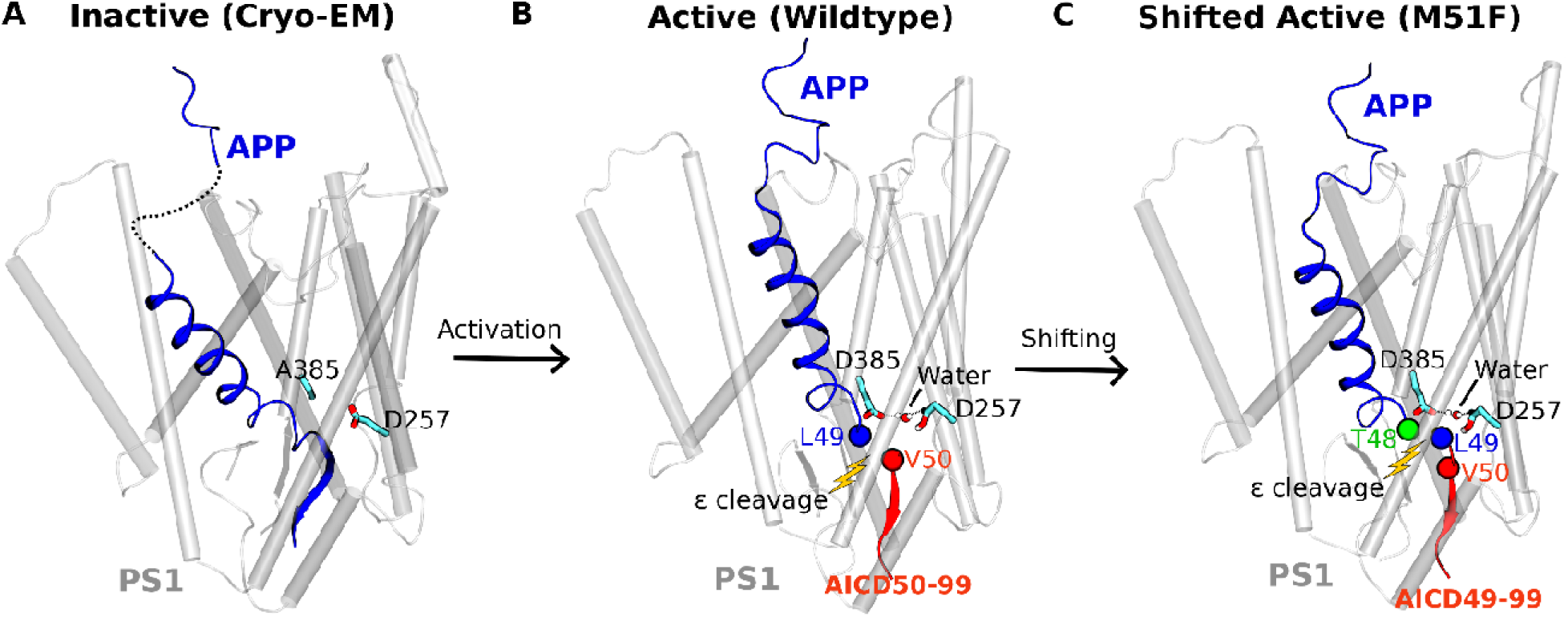
Summary of the (A) inactive cryo-EM, (B) active (wildtype) and (C) shifted active (M51F) conformational states of the APP substrate-bound γ-secretase. Distinct AICD products were generated from the wildtype and M51F mutant APP. The complementary simulations and experiments have revealed mechanisms of the γ-secretase activation and its ε cleavage of the APP substrate.

GaMD simulations on I45F and T48P APP substrate-bound γ-secretase revealed faster activation of PS1 for proteolysis at the ε cleavage site between Leu49-Val50 with these two FAD mutations compared to the complex with wildtype APP substrate. These observations were consistent with MS analysis of AICD proteolytic products: the two FAD-mutant substrates were cleaved by γ-secretase with a greater AICD50-99/AICD49-99 ratio than was the wildtype substrate. Moreover, the M51F mutation resulted in dramatic conformational changes of APP **(Figure 6C** and **Movie S3)**, setting up ε cleavage between Thr48-Leu49. These results were entirely consistent with the known incompatibility of Phe in the P2’ position.^10^ MS experimental results also showed the major AICD product generated by γ-secretase from the M51F-mutant APP substrate was due to cleavage between Thr48-Leu49. Little or no cleavage occurred between Leu49-Val50. In addition, western blotting revealed a substantial increase in the total AICD production in the *in-vitro* γ-secretase assay for the M51F-mutant APP substrate compared to the wildtype APP substrate. In contrast, I45F and T48P mutant APP-bound γ-secretase showed similar amount of AICD production as the wildtype APP bound γ-secretase. This was in exceptional agreement with the GaMD simulation: the low-energy inhibited state was observed in the free energy profiles of the wildtype, I45F and T48P mutant APP bound γ-secretase, but absent in the M51F mutant APP system. These strong correlations between the GaMD simulations and biochemical experiments provided substantial validity to our dynamic model of γ-secretase.

The active-site S1’, S2’ and S3’ subpockets were visualized in the wildtype active, I45F active, T48P active and M51F shifted active conformations of PS1 obtained from GaMD simulations. The protein residues **(Figure 5** and **Table S3)** found in the S1’ and S3’ subpockets of the active wildtype, I45F and T48P conformations were the same as those identified in a recent computational study by Hitzenberger et.al.^25^ However, the S2’ pocket of the wildtype active PS1 was identified in a distinct location that shifted by ∼4 Å towards the APP C-terminus from the previously described S2’ pocket.^25^ The subpocket described by Hitzenberger et.al.^25^, on the other hand, appeared to be the S2’ pocket in the I45F and T48P active conformations and the shifted S3’ subpocket for the M51F APP **(Figure 5E)**. Shift of the S2’ subpocket from the wildtype active conformation to the I45F and T48P active conformations resulted from the simultaneous movements of the APP C-terminus and PS1 TM7 N-terminus towards the PS1 TM6a in order to maintain the β-sheet structure of this domain in the GaMD simulations **(Figure S12)**. Therefore, the GaMD simulations revealed a newly identified S2’ subpocket for wildtype APP, while the previously described S2’ subpocket^25^ was used as the S2’ for I45F and T48P APP as well as the shifted S3’ for the M51F APP **(Figure 5)**.

In summary, we have combined all-atom GaMD simulations with MS and western blotting experiments to probe the mechanisms of γ-secretase activation and its ε cleavage of the wildtype and mutant APP substrates. Extensive GaMD simulations using the latest cryo-EM structures of γ-secretase have captured spontaneous activation of the enzyme, for which the active-site Asp385 has been restored and the artificial enzyme-substrate disulfide bond has been removed. The active conformation is characterized by water-bridged hydrogen bonds between the two catalytic aspartates, one of which formed another hydrogen bond with the carbonyl oxygen of the target scissile amide bond for the ε cleavage of APP. Free energy calculations of the GaMD simulations also allowed us to identify distinct intermediate, inhibited and shifted active conformational states of γ-secretase. The simulations predicted ε cleavage preferences of the wildtype and three mutants of APP that were highly consistent with MS and western blotting experimental findings of the AICD species. The validated GaMD simulations were then used to interpret the experimental data at an atomistic level. Remarkably, the M51F mutation shifted the ε cleavage site of APP from the amide bond between Leu49-Val50 to the Thr48-Leu49 bond, generating predominantly the AICD49-99 fragment instead of the AICD50-99 as detected by MS. Finally, the GaMD simulations have systematically revealed the active-site S1’, S2’ and S3’ subpockets of γ-secretase that interacted with the P1’, P2’ and P3’ residues in the wildtype and mutant APP. This provides an in-depth picture of the ε proteolytic cleavage of different APP substrates by γ-secretase. The GaMD method is apparently very well suited for the study of this extremely slow-acting membrane protease complex. In order to fully understand the functional mechanisms of γ-secretase, further simulation and experimental studies have been planned on the tripeptidase activity of the enzyme and effects of FAD mutations in both the APP substrate and γ-secretase. These studies are expected to greatly facilitate rational drug design targeting γ-secretase for the AD therapeutic treatments.

## Materials and Methods

### Cloning

All mutations in C100 FLAG were introduced by site-directed mutagenesis (QuickChange Lightning Site Directed Mutagenesis kit, Agilent) in pET 22b vector. All constructs were verified by sequencing by ACGT.

### C100-FLAG substrates purification

*E. coli* BL21 cells were grown in LB media until OD_600_ reached 0.6. Cells were induced with 0.5 mM IPTG and were grown post induction for 4 hours. The cells were then pelleted by centrifugation and resuspended in 50 mM HEPES pH 8, 1% Triton X-100. The cells were lysed by French press and the lysate was incubated with anti-FLAG M2-agarose beads from SIGMA. Bound substrates were then eluted from the beads with 100 mM Glycine pH 2.5, 0.25% NP-40 detergent and then neutralized with Tris HCl prior to being stored at −80°C.

### γ-secretase expression and purification

γ-secretase was expressed in HEK 395F cells by transfection with pMLINK vector containing all four components (Presenilin-1, Pen-2, Aph-1, Nicastrin) of γ-secretase complex (provided by Yigong Shi). For transfection, HEK 395F cells were grown in unsupplemented Freestyle 293 media (Life Technologies, 12338-018) until cell density reached 2×10^6^ cells/ml. 150 μg of vector was mixed with 450 μg of 25 kDa linear polyethylemimines (PEI) and incubated for 30 minutes at room temperature. The DNA-PEI mixtures were added to HEK cells and cells were grown for 60 hours. The cells were harvested, and γ-secretase was purified as described previously.^10^

### In vitro γ-secretase assay and detection of AICD species

γ -secretase purification and assays were carried out as described previously.^10^ Briefly, 30 nM purified γ-secretase was dissolved into total brain lipid extract (Avanti) in 50 mM HEPES pH 7.0, 150 mM NaCl, 0.25% CHAPSO. The detergent/lipid/enzyme solution was mixed with SM-2 bio-beads (Bio-Rad) for 2 h at 4 ^0^C to remove the detergent. After removal of the bio beads, the proteoliposome solution was mixed with 3 mM recombinant C100 substrates to initiate the cleavage reaction. The reaction was carried out for 16 h at 37 ^0^C. After 16 h, AICD-Flag products were isolated by immunoprecipitation with anti-FLAG M2 beads (SIGMA) in 10 mM MES pH 6.5, 10 mM NaCl, 0.05% DDM detergent overnight at 4 ^0^C. AICD products were then eluted from the anti-FLAG beads with acetonitrile:water (1:1) with 0.1% trifluoroacetic acid. The elutes were run on a Bruker MALDI-TOF mass spectrometer.

### Western blotting

Samples from γ-secretase assays were run on 4-12% bis-tris gel and transferred into PVDF membrane. The membrane was treated with 5% dry milk in PBS Tween-20 for 1 h at ambient temperature. The membrane was then incubated with the anti-Flag M2 antibodies at 4 °C overnight. The membrane was washed 3 times with PBS Tween-20 and was incubated with anti-mouse secondary antibodies for 1 h. The membrane was washed and imaged for chemiluminescence.

### Gaussian Accelerated Molecular Dynamics (GaMD)

GaMD is an enhanced sampling technique, in which a harmonic boost potential is added to smooth the potential energy surface and reduce the system energy barriers.^27^ GaMD is able to accelerate biomolecular simulations by orders of magnitude.^34, 54^ GaMD does not need predefined collective variables. Moreover, because GaMD boost potential follows a gaussian distribution, biomolecular free energy profiles can be properly recovered through cumulant expansion to the second order.^27^ GaMD has successfully overcome the energetic reweighting problem in free energy calculations that was encountered in the previous accelerated molecular dynamics (aMD)method^30, 32^ for free energy calculations of large molecules. GaMD has been implemented in widely used software packages including AMBER^27, 55^, NAMD^33^ and GENESIS^56^. A brief summary of GaMD is provided here.

Consider a system with *N* atoms at positions 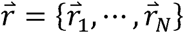 When the system potential 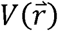 is lower than a reference energy *E*, the modified potential 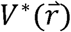 of the system is calculated as:

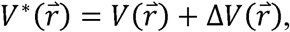

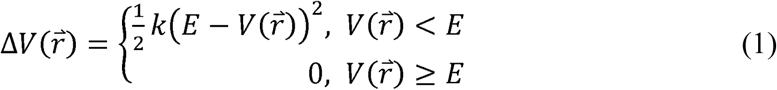

where *k* is the harmonic force constant. The two adjustable parameters *E* and *k* are automatically determined based on three enhanced sampling principles.^27^ The reference energy needs to be set in the following range:

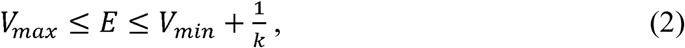

where *V*_*max*_ and *V*_*min*_ are the system minimum and maximum potential energies. To ensure that Eqn. (2) is valid, *k* has to satisfy: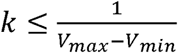 Let us define 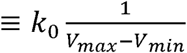, then 0 ≤ k_0_ ≤1

The standard deviation of Δ*v* needs to be small enough (i.e., narrow distribution) to ensure proper energetic reweighting^57^: σ_Δ*V*_ = *k* (*E _ V*_avg_)σ_*V*_ ≤ σ_0_ where *V*_avg_ and σ_*V*_ are the average and standard deviation of the system potential energies, σ_Δ*V*_ is the standard deviation of Δ*v* with σ_0_ as a user-specified upper limit (e.g., 10*k*_*B*_T) for proper reweighting. When *E* is set to the lower bound *E=V*_*max*_, *k*_0_ can be calculated as:

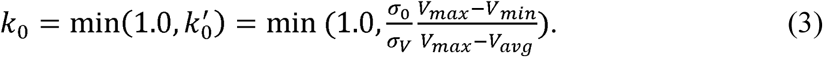

Alternatively, when the threshold energy *E* is set to its upper bound 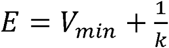, *k*_0_ is set to:

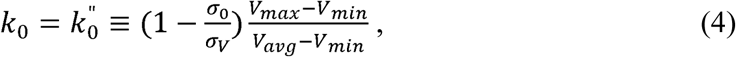

if 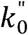 is found to be between *0* and *1*. Otherwise, *k*_0_ is calculated using Eqn. (3).

Similar to aMD, GaMD provides schemes to add only the total potential boost Δ*v*_*P*_, only dihedral potential boost Δ*v*_*D*_, or the dual potential boost (both Δ*v*_*P*_ and Δ*v*_*D*_). The dual-boost simulation generally provides higher acceleration than the other two types of simulations^58^. The simulation parameters comprise of the threshold energy E for applying boost potential and the effective harmonic force constants, *k*_0*P*_ and *k*_0*D*_ for the total and dihedral potential boost, respectively.

### Energetic Reweighting of GaMD Simulations

To calculate potential of mean force (PMF)^59^ from GaMD simulations, the probability distribution along a reaction coordinate is written as *p* * (A)Given the boost potential 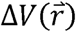 of each frame, *p* *(A)can be reweighted to recover the canonical ensemble distribution, *p* (A), as:

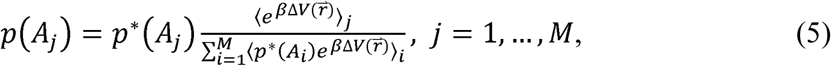

where *M* is the number of bins, *β*,= *k*_*B*_*T* and 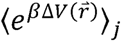 is the ensemble-averaged Boltzmann factor of 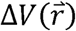 for simulation frames found in the *j*^th^ bin. The ensemble-averaged reweighting mfactor can be approximated using cumulant expansion:

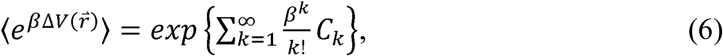

where the first two cumulants are given by

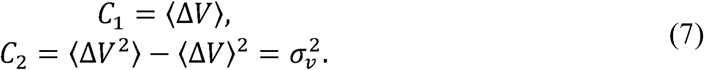

The boost potential obtained from GaMD simulations usually follows near-Gaussian distribution. Cumulant expansion to the second order thus provides a good approximation for computing the reweighting factor^27, 57^. The reweighted free energy *F (A) =* − *K*_*B*_*T* ln *P (A)* is calculated as:

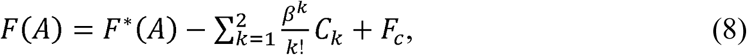

where *F*\**(A) =* − *K*_*B*_*T* In *P*\**(A)* is the modified free energy obtained from GaMD simulation and *F*_*c*_ is a constant.

### System Setup

The earlier published cryo-EM structure of γ-secretase bound by Notch (PDB: 6IDF)^53^ was used for initial GaMD simulations. This system was used to optimize our simulation protocol, especially the protonation state of aspartates in the active site. Another cryo-EM structure of γ-secretase bound by APP (PDB: 6IYC)^12^ was used to perform further GaMD simulations as per the optimized protocol. For the wildtype enzyme, residue Asp385 that was mutated to Ala at the active site in the cryo-EM structure was restored for setting up the simulation system. Similarly, the disulfide bond between Cys112 of PS1-Q112C and Cys24 of APP-V24C were removed, and the wildtype residues were restored for simulation setup. Five unresolved residues at the N-terminus of APP substrate C83 were added through homology modelling using SWISS-MODEL.^60^ All chain termini were capped with neutral groups, i.e. the acetyl group (ACE) for the N-terminus and methyl amide group (CT3) for C terminus. Protein residues were set to the standard CHARMM protonation states at neutral pH with the *psfgen* plugin in VMD.^61^ Then the complex was embedded in a 1-palmitoyl-2-oleoyl-sn-glycero-3-phosphocholine (POPC) bilayer with all overlapping lipid molecules removed using the *Membrane* plugin in VMD^61^ **(Figure S1)**. The system charges were then neutralized at 0.15 M NaCl using the *Solvate* plugin in VMD.^61^ Periodic boundary conditions were applied on the simulation systems. The simulation systems of γ-secretase bound by APP are summarized in **Table 1**.

For APP-mutant simulations systems, isoleucine, threonine and methionine were mutated to phenylalanine, proline and phenylalanine computationally at the 29^th^, 32^nd^ and 35^th^ residue of APP substrate, respectively. These corresponded to I45F, T48P and M51F mutations as per the numbering based on C99, the substrate that was cleaved to Aβ, although the actual substrate in the model was C83.

### Simulation Protocol

The CHARMM36 parameter set^62^ was used for the protein and POPC lipids. Initial energy minimization and thermalization of the γ-secretase complex followed the same protocol as used in the previous GaMD simulations of membrane proteins.^34, 63^ The simulation proceeded with equilibration of lipid tails. With all the other atoms fixed, the lipid tails were energy minimized for 1000 steps using the conjugate gradient algorithm and melted with constant number, volume, and temperature (NVT) run for 0.5 ns at 310 K. Each system was further equilibrated using constant number, pressure, and temperature (NPT) run at 1 atm and 310 °K for 10 ns with 5 kcal (mol Å^2^)^-1^ harmonic position restraints applied to the protein. Further equilibration of the systems was performed using an NPT run at 1 atm and 310 °K for 0.5 ns with all atoms unrestrained. Conventional MD simulation was performed on each system for 10 ns at 1 atm pressure and 310 °K with a constant ratio constraint applied on the lipid bilayer in the X-Y plane. The GaMD simulations were carried out using AMBER 18.^27, 55^ Dual-boost GaMD simulations were performed to study the substrate-bound γ-secretase complex (**Table 1**). In the GaMD simulations, the threshold energy *E* for adding boost potential was set to the lower bound, i.e. *E = V*_max_.^27, 33^ The simulations included 50 ns equilibration after adding the boost potential and then multiple independent production runs lasting 1-2 μs with randomized initial atomic velocities. GaMD production simulation frames were saved every 0.2 ps for analysis.

### Simulation analysis

The VMD^61^ and CPPTRAJ^64^ tools were used for trajectory analysis. In particular, distance was calculated between the Cγ atoms of catalytic aspartate residues. Hydrogen bond distance was calculated between donor protonated oxygen atom of PS1 Asp257 and the acceptor carbonyl oxygen atom of APP substrate residue Leu49, Thr48 or Ile47. Root-mean-square fluctuations (RMSFs) were calculated for the protein residues, averaged over three independent GaMD simulations and color coded for schematic representation of each complex system. The CPPTRAJ was used to calculate the protein secondary structure plots. The *PyReweighting* toolkit^57^ was applied to reweight GaMD simulations for free energy calculations by combining all simulation trajectories for each system. A bin size of 1 Å was used for the PMF calculation of distances. The cutoff was set to 500 frames in each bin for calculating the 2D PMF profiles. Protein snapshots were taken every 1 ps for structural clustering. Clustering was performed on the GaMD simulations of wildtype, I45F, T48P and M51F-mutant APP bound γ-secretase based on the RMSD of PS1 using hierarchical agglomerative algorithm in CPPTRAJ^64^ generating ∼10 representative structural clusters for each system. The top structural cluster was identified as the representative active (wildtype) and shifted active conformational states of the wildtype and M51F mutant APP bound γ-secretase systems, respectively. The top structural cluster was also identified as the active (I45F and T48P) conformational state of the I45F and T48P mutant APP bound γ-secretase.

## ASSOCIATED CONTENT

### Supporting Information

Supplementary file containing Supporting Methods and Materials, **Figures S1 - S12** and **Table S1**. The Supporting Information is available free of charge on the ACS Publications website.

#### Acknowledgements

This work used supercomputing resources with allocation award TG-MCB180049 through the Extreme Science and Engineering Discovery Environment (XSEDE), which is supported by National Science Foundation grant number ACI-1548562, and project M2874 through the National Energy Research Scientific Computing Center (NERSC), which is a U.S. Department of Energy Office of Science User Facility operated under Contract No. DE-AC02-05CH11231, and the Research Computing Cluster at the University of Kansas. This work was supported in part by the startup funding in the College of Liberal Arts and Sciences at the University of Kansas (Y.M.) and GM122894 from the National Institutes of Health (M.S.W.).

## Table of Contents/Abstract Graphics

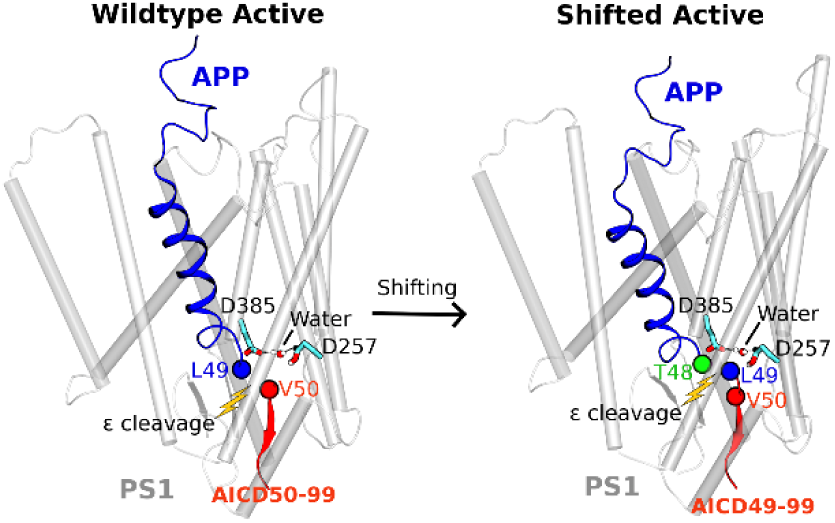

Complementary accelerated molecular simulations, mass spectrometry and western blotting experiments have revealed spontaneous activation of γ-secretase bound by the wildtype amyloid precursor protein (APP) substrate and shift of the ε cleavage site in processing of mutant APP, yielding different APP intracellular domains (AICD50-99 and AICD49-99).

