## Supporting Information for "Mechanisms of γ-Secretase Activation and Substrate Processing"

**Table of Contents:**

1. **Supporting Methods and Materials**
2. **Supporting Figures (Figures S1 - S12)**
3. **Supporting Tables (Table S1)**
4. **References**

### 1. Supporting Methods and Materials

#### Cloning

All mutations in C100 FLAG were introduced by site-directed mutagenesis (QuickChange Lightning Site Directed Mutagenesis kit, Agilent) in pET 22b vector. All constructs were verified by sequencing by ACGT.

**Gaussian Accelerated Molecular Dynamics (GaMD)**

GaMD is an enhanced sampling technique, in which a harmonic boost potential is added to smooth the potential energy surface and reduce the system energy barriers.[2] GaMD is able to accelerate biomolecular simulations by orders of magnitude.[3-4] GaMD does not need predefined collective variables. Moreover, because GaMD boost potential follows a gaussian distribution, biomolecular free energy profiles can be properly recovered through cumulant expansion to the second order.[2] GaMD has successfully overcome the energetic reweighting problem in free energy calculations that was encountered in the previous accelerated molecular dynamics (aMD)method[5-6] for free energy calculations of large molecules. GaMD has been implemented in widely used software packages including AMBER[2, 7], NAMD[8] and GENESIS[9]. A brief summary of GaMD is provided here.

Consider a system with $N$ atoms at positions $\vec{r} = \{\vec{r}_1, \cdots, \vec{r}_N\}$. When the system potential $V(\vec{r})$ is lower than a reference energy $E$, the modified potential $V^*(\vec{r})$ of the system is calculated as:

$$V^*(\vec{r}) = V(\vec{r}) + \Delta V(\vec{r}),$$

$$\Delta V(\vec{r}) = \begin{cases} \frac{1}{2} k\left(E - V(\vec{r})\right)^2, & V(\vec{r}) < E \\ 0, & V(\vec{r}) \geq E \end{cases} \quad (1)$$

where $k$ is the harmonic force constant. The two adjustable parameters $E$ and $k$ are automatically determined based on three enhanced sampling principles.[2] The reference energy needs to be set in the following range:

$$V_{max} \leq E \leq V_{min} + \frac{1}{k}, \quad (2)$$

where $V_{max}$ and $V_{min}$ are the system minimum and maximum potential energies. To ensure that Eqn. (2) is valid, $k$ has to satisfy: $k \leq \frac{1}{V_{max} - V_{min}}$ Let us define $\equiv k_0 \frac{1}{V_{max} - V_{min}}$, then $0 < k_0 \leq 1$. The standard deviation of $\Delta V$ needs to be small enough (i.e., narrow distribution) to ensure proper energetic reweighting[10]: $\sigma_{\Delta V} = k\left(E - V_{avg}\right)\sigma_V \leq \sigma_0$ where $V_{avg}$ and $\sigma_V$ are the average and standard deviation of the system potential energies, $\sigma_{\Delta V}$ is the standard deviation of $\Delta V$ with $\sigma_0$ as a user-specified upper limit (e.g., 10$k_B$T) for proper reweighting. When $E$ is set to the lower bound $E=V_{max}$, $k_0$ can be calculated as:

$$k_0 = \min(1.0, k_0') = \min\left(1.0, \frac{\sigma_0}{\sigma_V}\frac{V_{max}-V_{min}}{V_{max}-V_{avg}}\right). \tag{3}$$

Alternatively, when the threshold energy $E$ is set to its upper bound $E = V_{min} + \frac{1}{k}$, $k_0$ is set to:

$$k_0 = k_0'' \equiv \left(1 - \frac{\sigma_0}{\sigma_V}\right)\frac{V_{max}-V_{min}}{V_{avg}-V_{min}}, \tag{4}$$

if $k_0''$ is found to be between *0* and *1*. Otherwise, $k_0$ is calculated using Eqn. (3).

Similar to aMD, GaMD provides schemes to add only the total potential boost $\Delta V_P$, only dihedral potential boost $\Delta V_D$, or the dual potential boost (both $\Delta V_P$ and $\Delta V_D$). The dual-boost simulation generally provides higher acceleration than the other two types of simulations[11]. The simulation parameters comprise of the threshold energy E for applying boost potential and the effective harmonic force constants, $k_{0P}$ and $k_{0D}$ for the total and dihedral potential boost, respectively.

**Energetic Reweighting of GaMD Simulations**

To calculate potential of mean force (PMF)[12] from GaMD simulations, the probability distribution along a reaction coordinate is written as $p^*(A)$ . Given the boost potential $\Delta V(\vec{r})$ of each frame, $p^*(A)$ can be reweighted to recover the canonical ensemble distribution, $p(A)$, as:

$$p(A_j) = p^*(A_j)\frac{\langle e^{\beta\Delta V(\vec{r})}\rangle_j}{\sum_{i=1}^{M}\langle p^*(A_i)e^{\beta\Delta V(\vec{r})}\rangle_i},\ j = 1, \ldots, M, \tag{5}$$

where $M$ is the number of bins, $\beta = k_B T$ and $\langle e^{\beta \Delta V(\vec{r})} \rangle_j$ is the ensemble-averaged Boltzmann factor of $\Delta V(\vec{r})$ for simulation frames found in the $j^{\text{th}}$ bin. The ensemble-averaged reweighting factor can be approximated using cumulant expansion:

$$\langle e^{\beta \Delta V(\vec{r})} \rangle = exp\left\{\sum_{k=1}^{\infty} \frac{\beta^k}{k!} C_k\right\}, \tag{6}$$

where the first two cumulants are given by

$$\begin{aligned} C_1 &= \langle \Delta V \rangle, \\ C_2 &= \langle \Delta V^2 \rangle - \langle \Delta V \rangle^2 = \sigma_v^2. \end{aligned} \tag{7}$$

**Restoration of wildtype γ-secretase for molecular dynamics simulations**

We performed initial GaMD simulations on the earlier published cryo-EM structure of γ-secretase bound by the Notch substrate (PDB: 6IDF[13]). Residue Ala385 at the active site was mutated back to aspartate, whereas the disulfide bond between the N-terminus of Notch substrate and hydrophobic loop 1 (HL1) loop was kept intact. In aspartyl proteases, proximity between the two active site Asp residues necessitates protonation of one of them, preventing charge repulsion. Testing GaMD simulations were performed to determine which of the two Asp residues was protonated in γ-secretase. We performed multiple 300 ns GaMD simulations **(Table 1)** on three different systems: the original cryo-EM structure, Asp257 protonated, and with Asp385 protonated. The distance time course plots **(Figure S2)** revealed that the system with protonated Asp257 in the N-terminal fragment (NTF) subunit of PS1 facilitated the activation of γ-secretase, as the two active-site aspartates approached each other to a distance of ~6-7 Å between the Cγ atoms. In contrast, simulations of the original cryo-EM structure did not show significant change from the starting Asp257:Cγ-Ala385:Cβ distance of ~10-11 Å. In the system with protonated Asp385 in the C-terminal fragment (CTF) of PS1, the two aspartates maintained a distance of ~10-

11 Å between the Cγ atoms of the two aspartates. Therefore, Asp257 was protonated in subsequent GaMD simulations being similar to the setup of a previous computational study[20].

Next, we proceeded to simulate γ-secretase bound by APP (PDB: 6IYC[14]). Residue Ala385 was similarly mutated back to Asp385 in the wildtype γ-secretase, which was compared to the original cryo-EM system in 300 ns GaMD simulations **(Table 1)**. The disulfide bond between substrate and enzyme was still kept. Free energy calculations showed that the active-site residues Asp257 and Ala385 maintained ~10 Å distance between their sidechain terminal C atoms in the cryo-EM system even though water molecules were observed entering the active site **(Figure S3A)**. The substrate remained distant from the active site residues, with ~5-6 Å between the Cγ atom of protonated Asp257 and the carbonyl oxygen of Leu49 in APP. In contrast, activation of γ-secretase was observed during three independent 300 ns GaMD simulations of the computationally restored wildtype enzyme **(Figure S3B, Movie S1)**. The protonated Asp257 formed a hydrogen bond with the carbonyl oxygen in Leu49 of the scissile amide bond in APP. Water molecules entered the PS1 active site. One water molecule was trapped between the two catalytic Asp residues through stable hydrogen bonds. This would induce nucleophilic attack of the carbonyl carbon of Leu49 by the activated water molecule, which is a key step for substrate proteolysis. The enzyme active site was thus well poised for proteolysis of APP for the ε cleavage between residues Leu49 and Val50. The two aspartates were ~7 Å apart between their Cγ atoms **(Figure S3B)**. The distance between the carbonyl carbon of Leu49 and the water oxygen was ~3.8 Å. Therefore, our GaMD simulations successfully captured activation of the APP-bound γ-secretase in the presence of the enzyme-substrate disulfide bond. Next, the artificial disulfide bond between the N-terminus of APP substrate and the HL1 loop of PS1 was also removed by computationally restoring the wildtype residues for further simulations as summarized in **Table 1.**

2. **Supporting Figures**

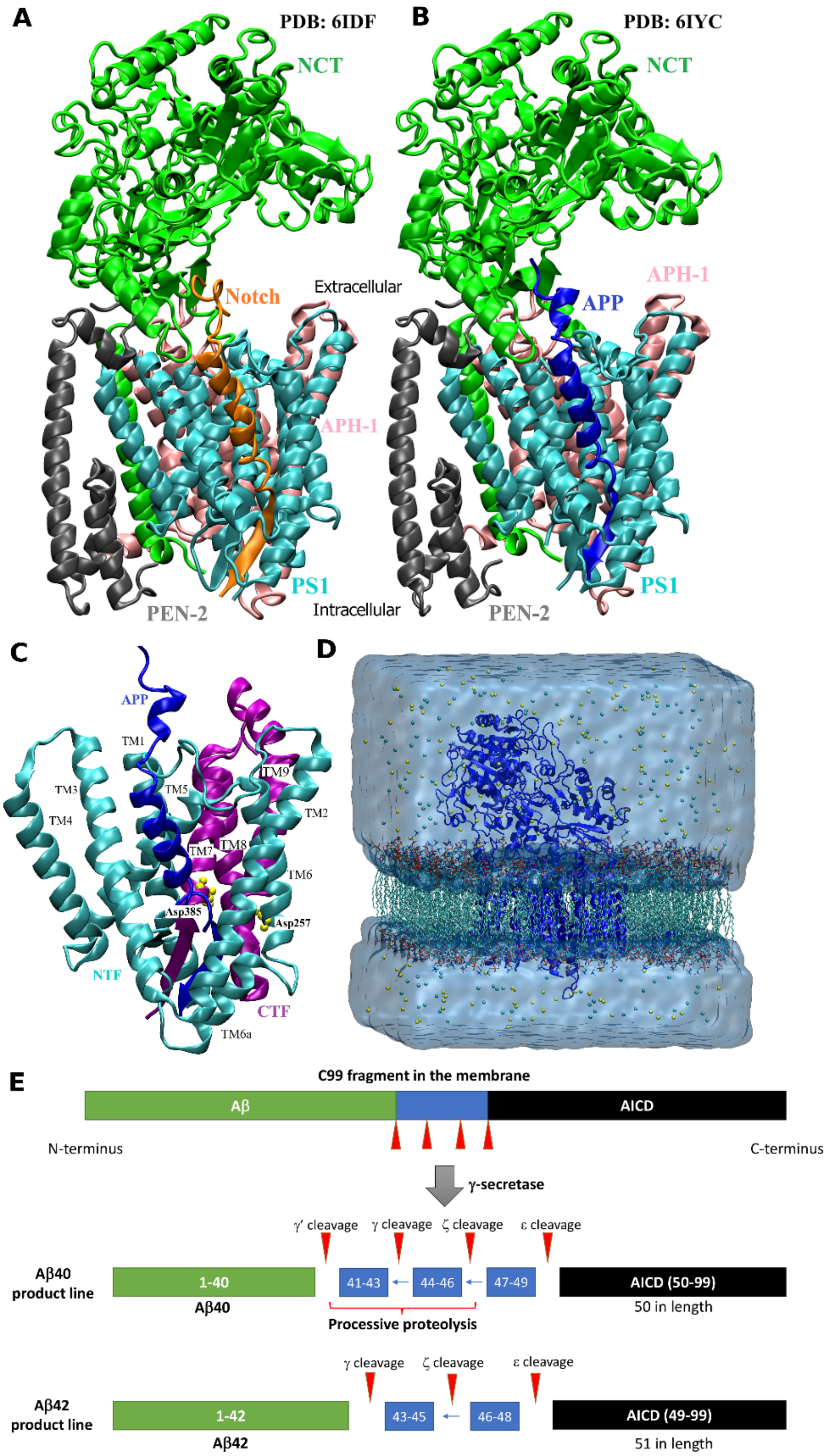


**Figure S1:** Ribbon representations of the (A) Notch- and (B) APP-bound γ-secretase complexes that include the presenilin (PS1), presenilin enhancer 2 (PEN2), anterior pharynx-defective 1

(APH1) and Nicastrin (NCT) subunits. (C) Representation of the catalytic PS1 domain of APP-bound γ-secretase. The transmembrane (TM) helices and active-site Asp385 and Asp257 residues are labelled. The N-terminal fragment (NTF) is colored in cyan and C-terminal fragment (CTF) is colored in purple (D) Computational model of γ-secretase complex in GaMD simulations. The protein was embedded into a POPC lipid bilayer and solvated in an aqueous medium of 0.15 M NaCl. (E) Schematic representation of APP substrate processing by γ-secretase.

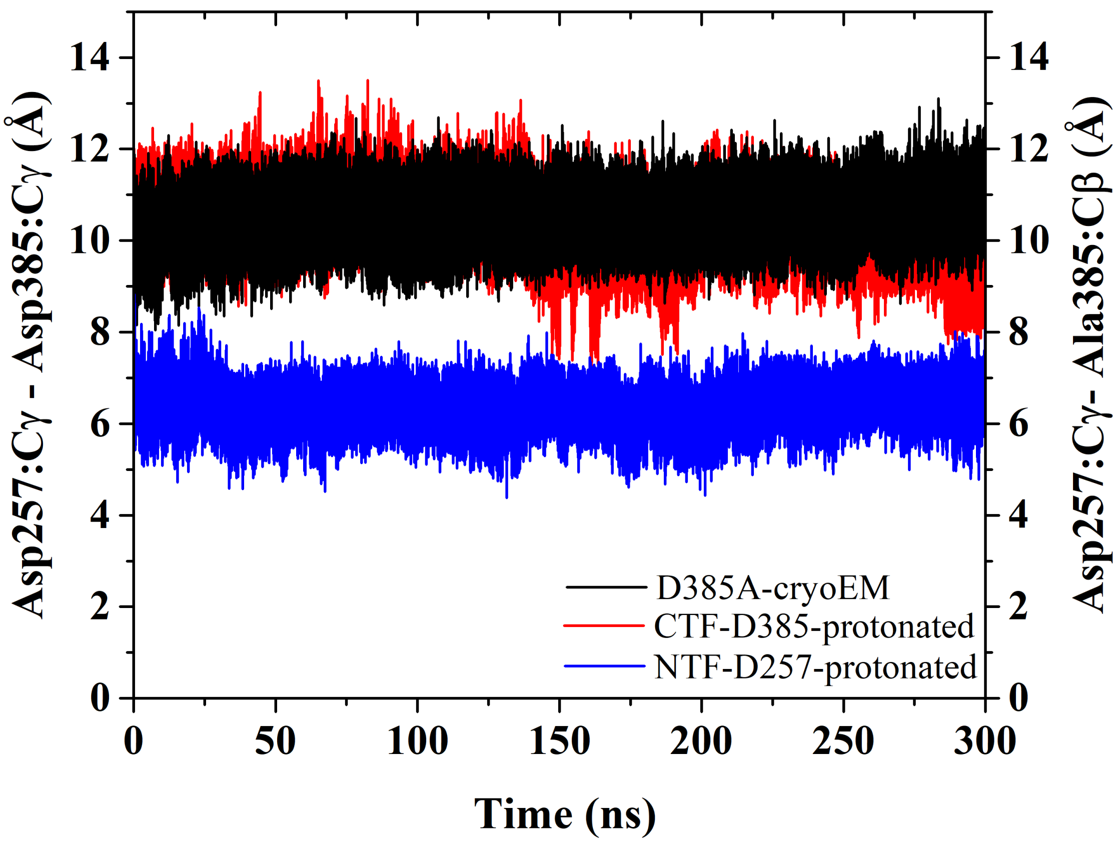


**Figure S2**: Time course of the Asp257:Cγ - Ala385:Cβ distance calculated from GaMD simulations of the cryo-EM D385A system (black) and Asp257:Cγ - Asp385:Cγ distance calculated from GaMD simulations of D385-protonated (red) and D257-protonated (blue) systems of Notch-bound γ-secretase complex. The disulfide bond between the N-terminus of Notch and PS1 HL1 loop was kept in these simulations.

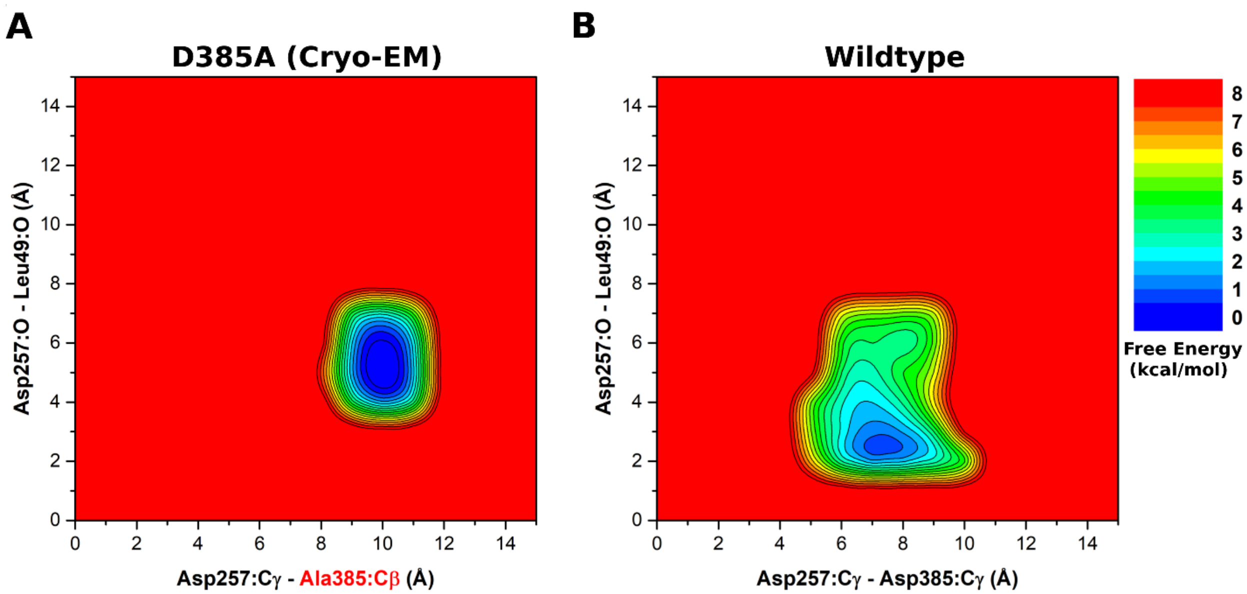


**Figure S3**: (A) 2D free energy profile of the Asp257:Cγ - Ala385:Cβ and Asp257:sidechain O - Leu49:O distances calculated from GaMD simulations of the D385A cryo-EM structure of γ-secretase bound by APP. (B) 2D free energy profile of the Asp257:Cγ - Asp385:Cγ and Asp257:p O - Leu49:protonated O distances calculated from GaMD simulations of the wildtype γ-secretase. The disulfide bond between the N-terminus of APP and PS1 HL1 loop was kept in these simulations.

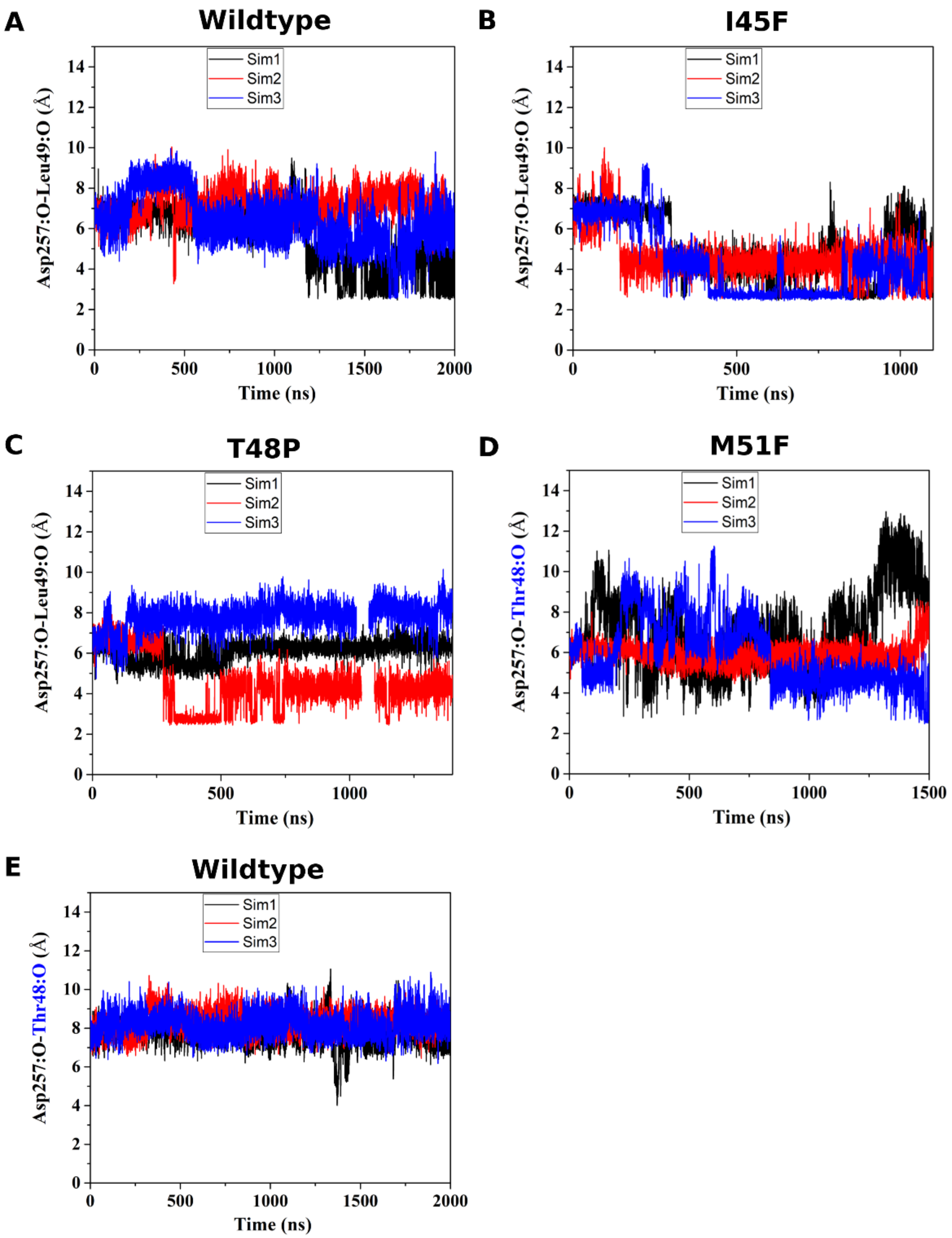


**Figure S4:** Time courses of the Asp257:protonated O - Leu49:O distance calculated from GaMD simulations of (A) wildtype, (B) I45F, and (C) T48P APP bound γ-secretase, and the Asp257:protonated O – Thr48:O distance calculated from GaMD simulations of the (D) M51F and (E) wildtype APP bound γ-secretase.

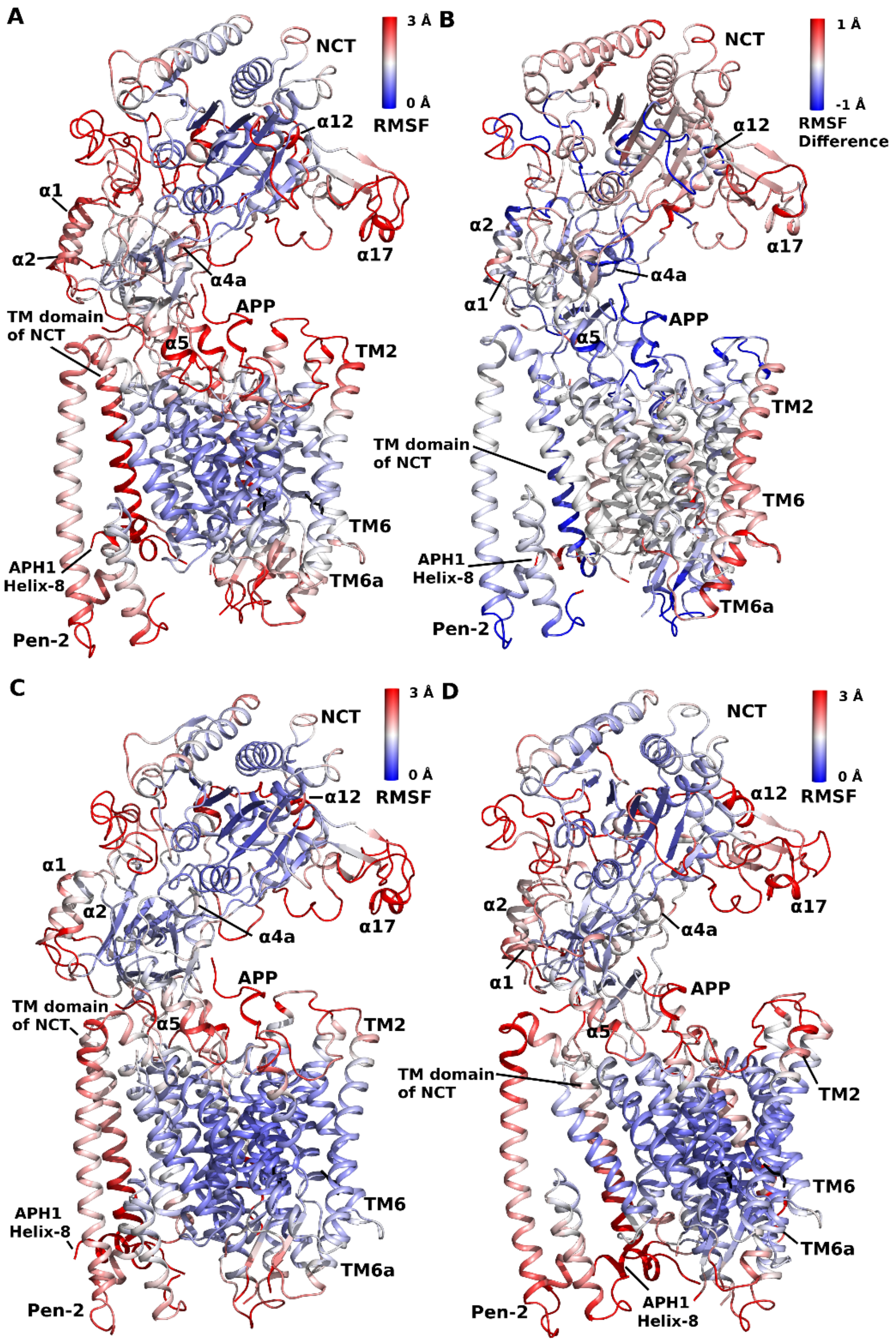


**Figure S5**: Comparison of structural flexibility of wildtype and mutant APP-bound γ-secretase calculated from GaMD simulations. (A) Root-mean-square fluctuations (RMSFs) of the wildtype APP-bound γ-secretase, (B) RMSF difference between the M51F and wildtype APP bound γ-secretase, (C) RMSFs of the I45F APP bound γ-secretase and (D) RMSFs of the T48P mutant APP-bound γ-secretase.

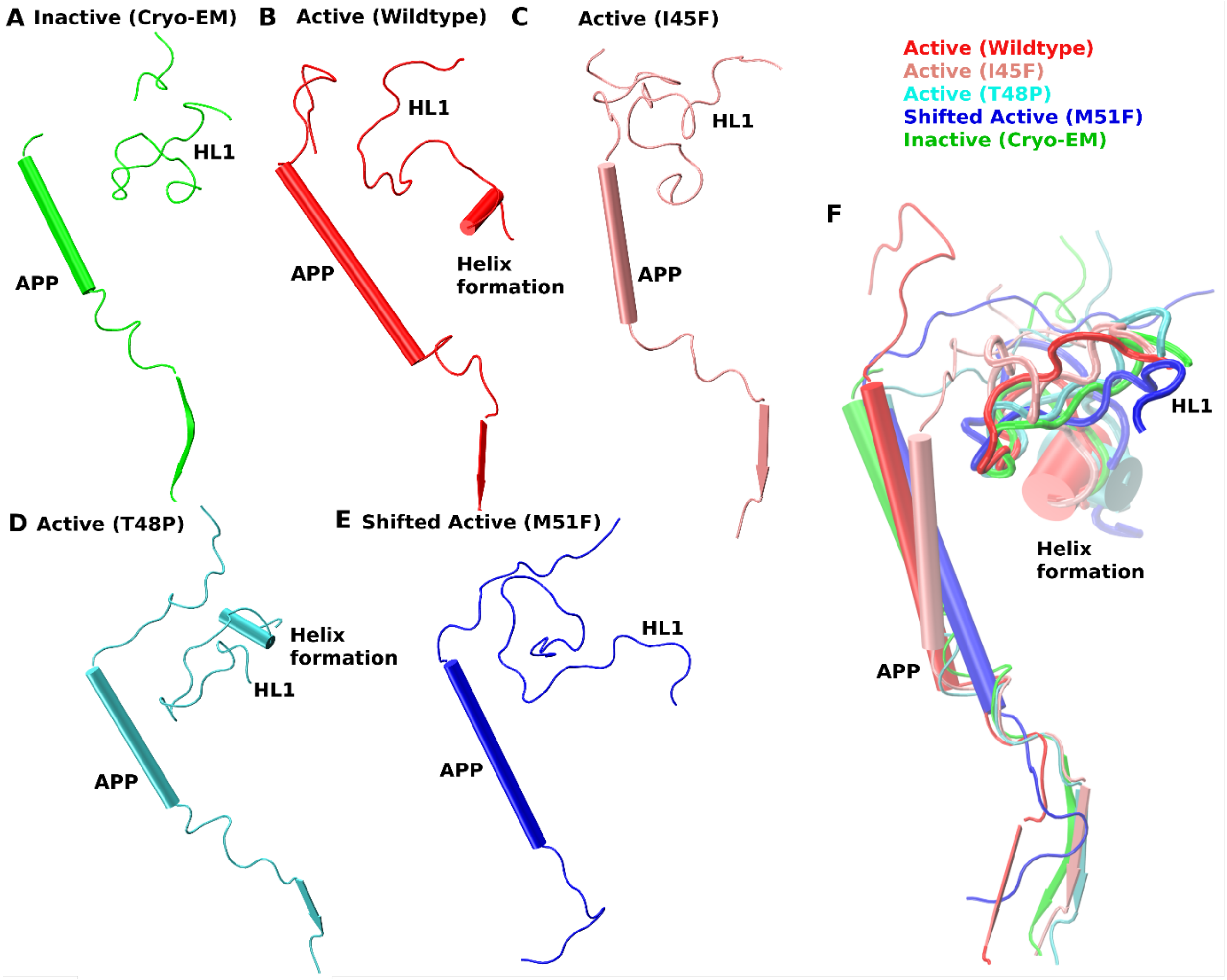


**Figure S6:** Side view of APP substrate and PS1 HL1 loop in the (A) inactive, representative active conformations of the (B) wildtype, (C) I45F and (D) T48P mutant APP-bound γ-secretase and the (E) Shifted conformation of M51F mutant APP-bound γ-secretase complex obtained from the GaMD simulations. (F) Comparison of APP substrate and PS1 HL1 loop in the different systems of APP-bound γ-secretase complex.

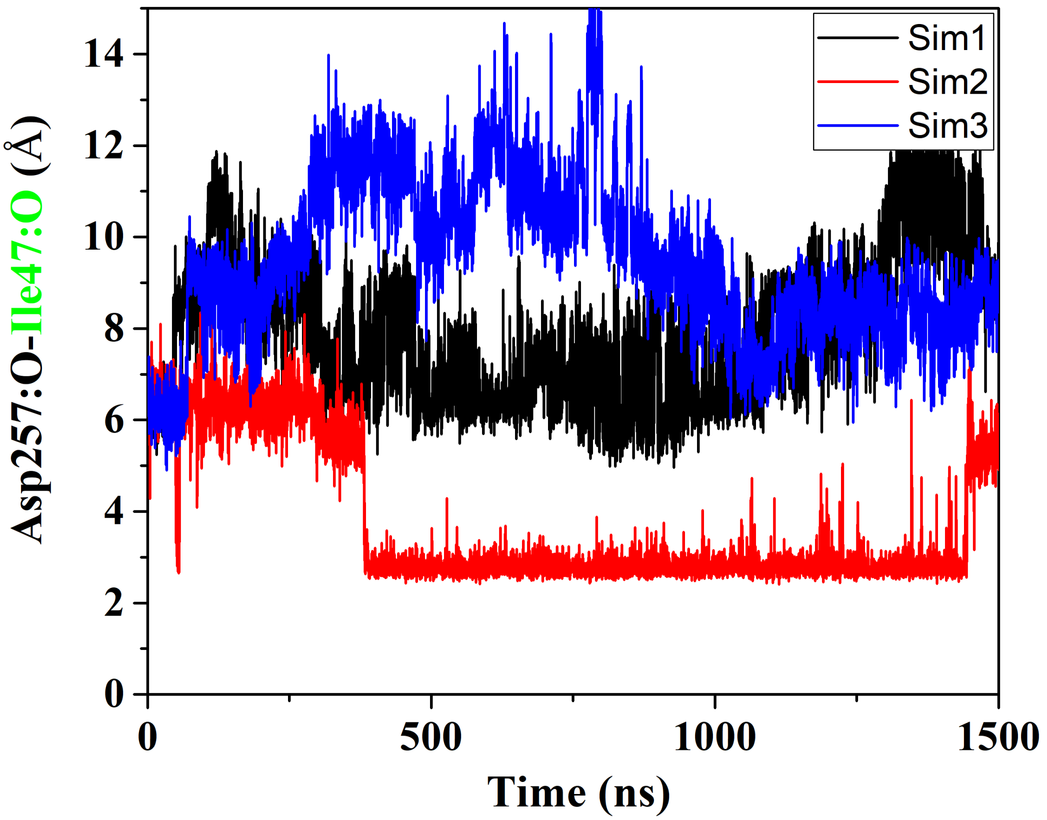


**Figure S7:** Time course of the Asp257:protonated O - Ile47:O distance calculated from GaMD simulations of the M51F APP bound γ-secretase.

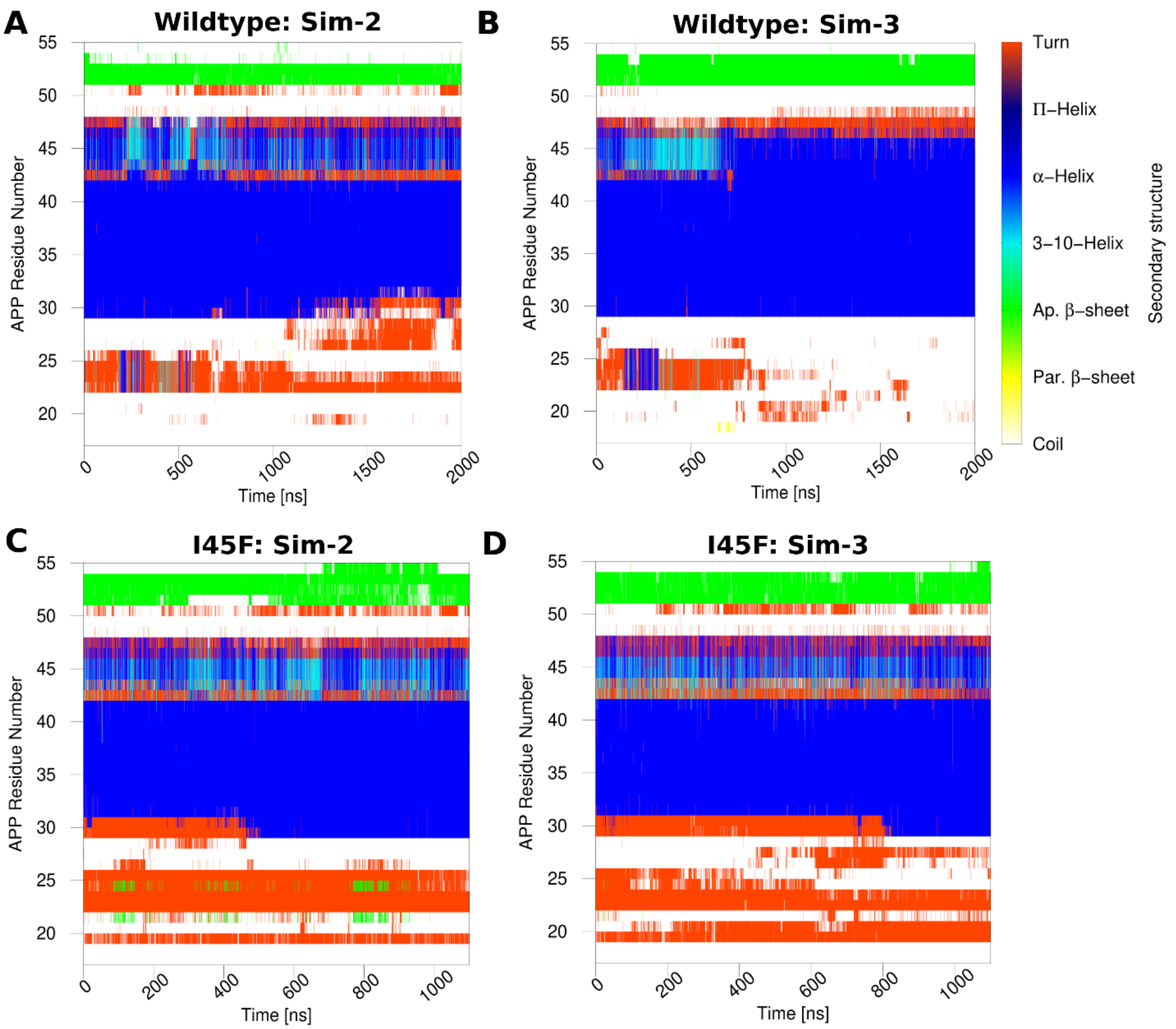


**Figure S8:** Time courses of the APP secondary structures in the wildtype and I45F forms as bound to γ-secretase calculated from GaMD simulations: (A) Sim-2 and (B) and Sim-3 for the wildtype (Sim-1 in Figure 4A), and (C) Sim-2 and (D) Sim-3 for the I45F mutant (Sim-1 in Figure 4B).

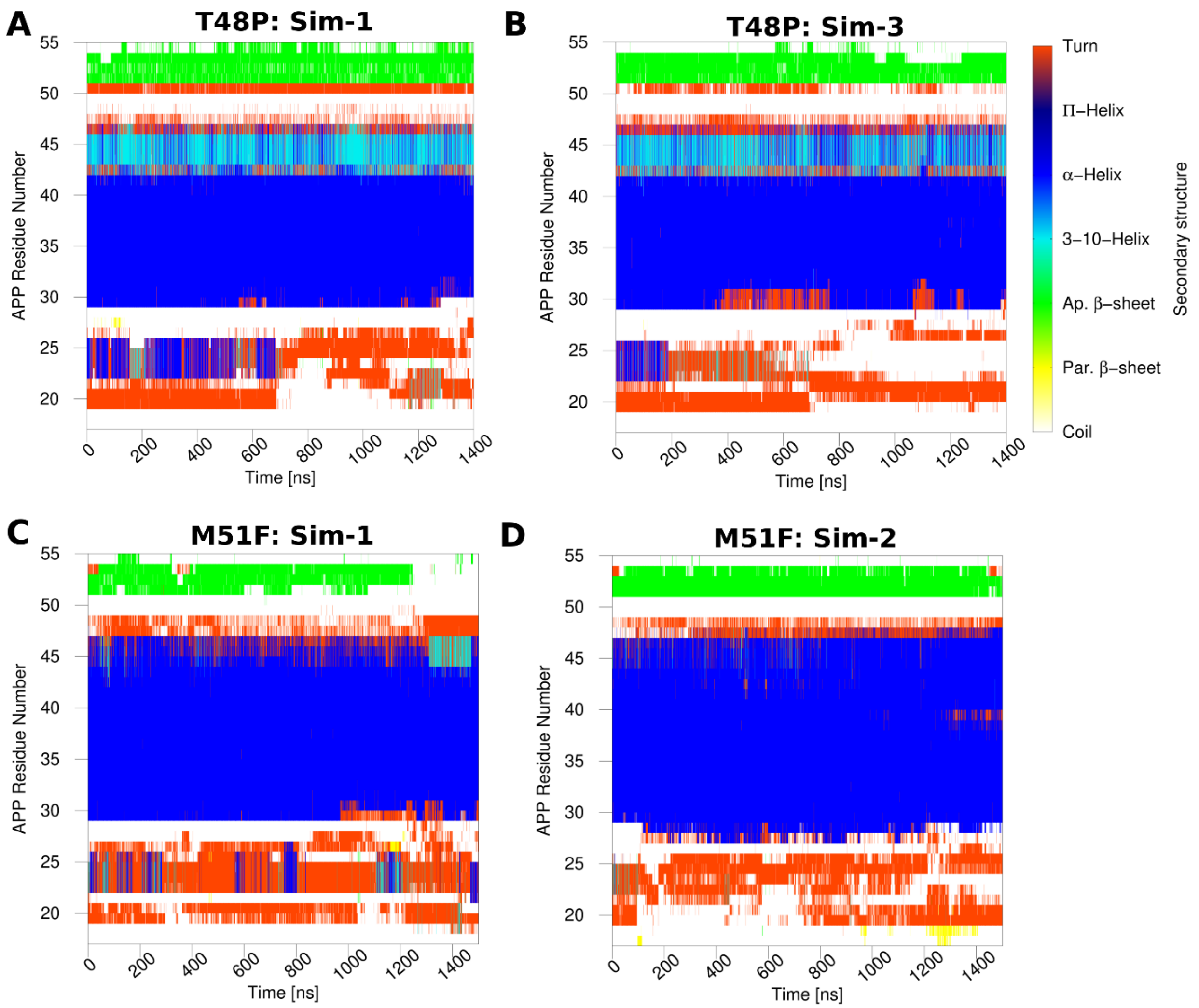


**Figure S9:** Time courses of the APP secondary structures in the T48P and M51F forms as bound to γ-secretase calculated from GaMD simulations: (A) Sim-1 and (B) and Sim-3 for the T48P mutant (Sim-2 in Figure 4C), and (C) Sim-1 and (D) Sim-2 for the M51F mutant (Sim-3 in 4D).

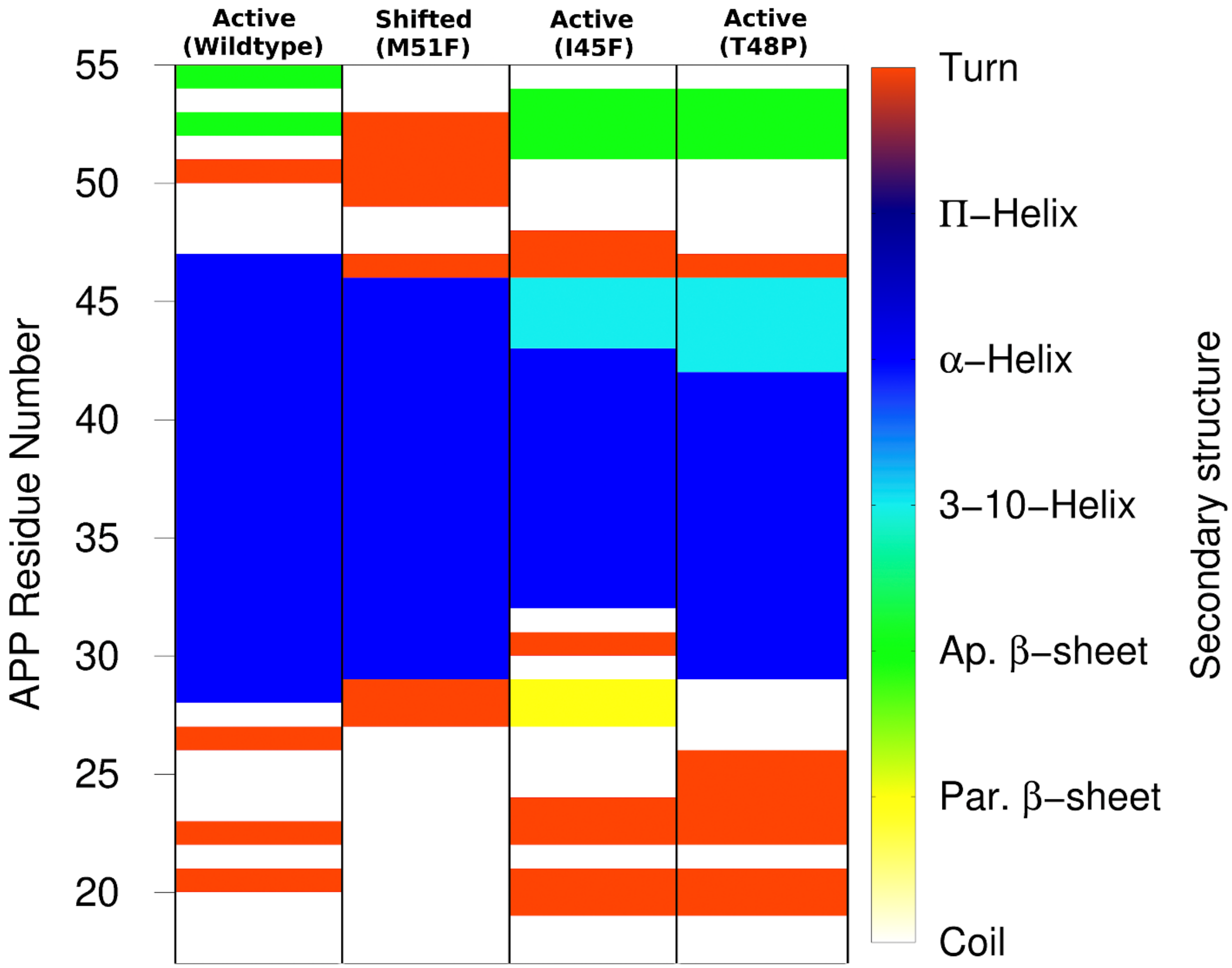


**Figure S10:** Secondary structures of the APP substrate in the representative active conformations of the wildtype, I45F and T48P mutant APP-bound γ-secretase and the shifted active conformation of M51F mutant APP-bound γ-secretase using the top ranked PS1-APP structural clusters obtained from the corresponding GaMD simulations.

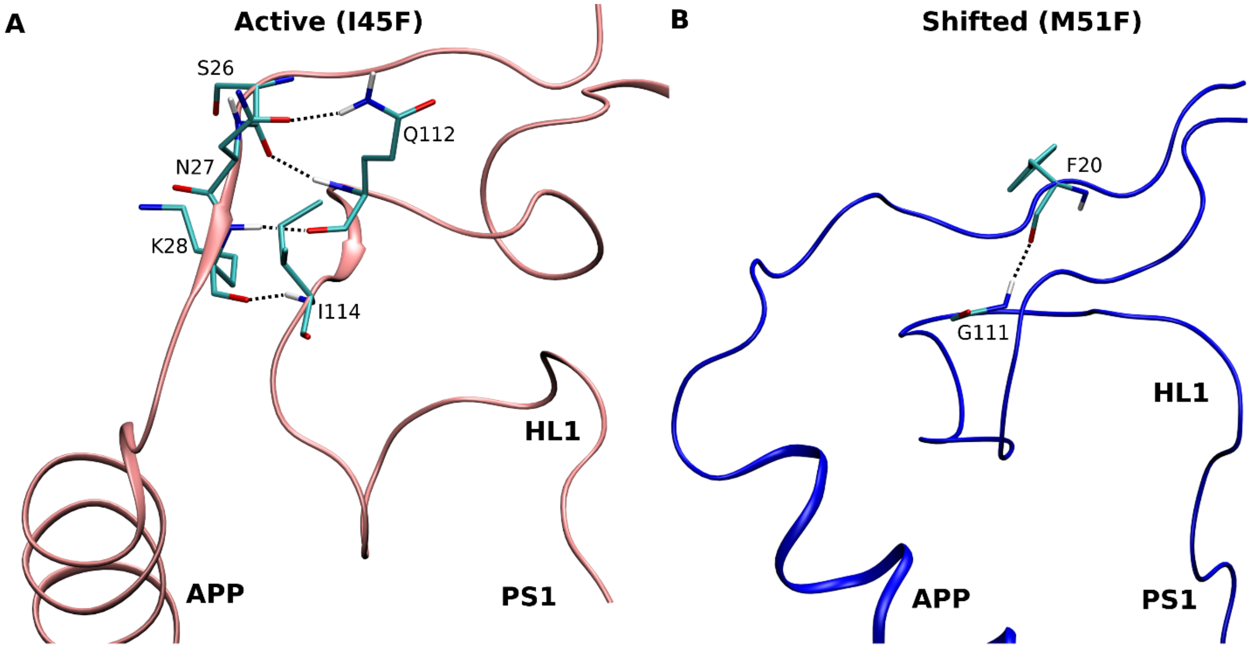


**Figure S11:** Hydrogen bonds formed between residues G111, Q112 and I114 of the PS1 HL1 loop and the N-terminus of APP substrate in the (A) Active and (B) Shifted conformations of γ-secretase obtained from the GaMD simulations of the I45F and M51F mutant APP systems.

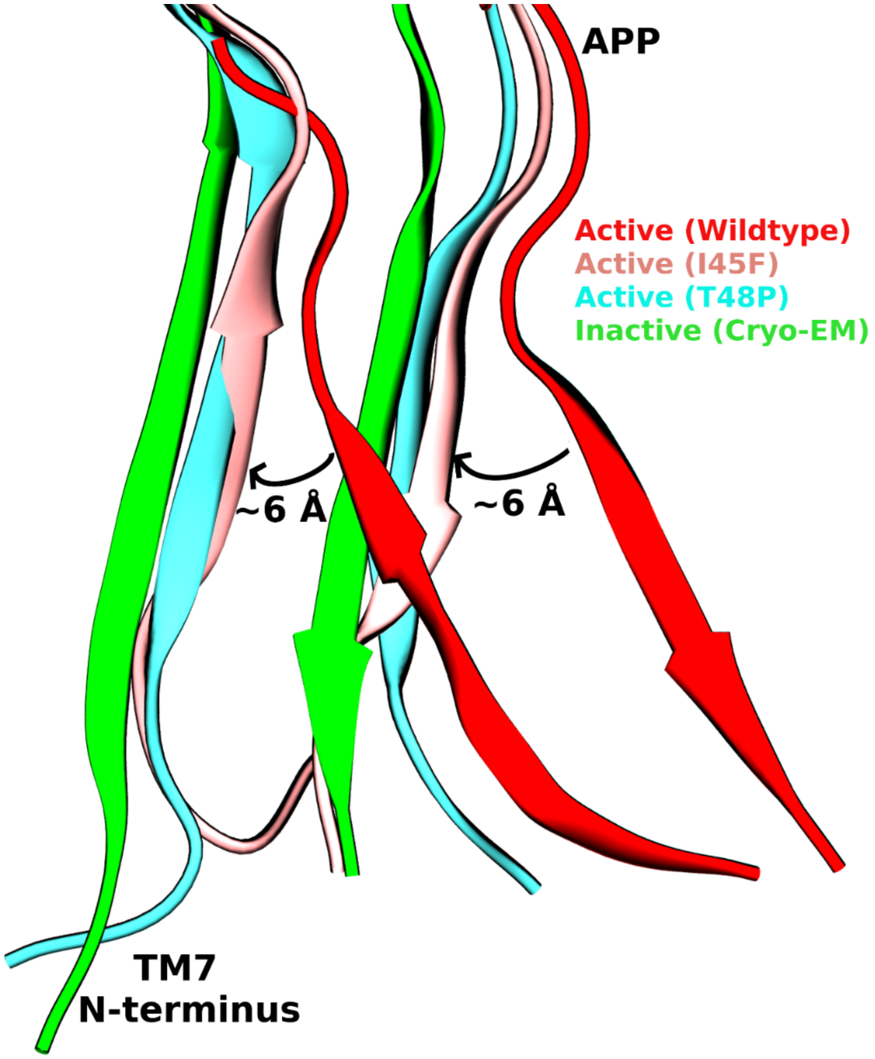


**Figure S12:** Comparison of the β-sheet conformational changes in the N-terminus of PS1 TM7 and the C-terminus of APP substrate in the inactive (cryo-EM) and active conformations of the wildtype, I45F and T48P mutant APP-bound γ-secretase observed in the GaMD simulations. Relative to the wildtype active conformation, the N-terminus of PS1 TM7 and the C-terminus of APP substrate moved towards the PS1 TM6a by ~6 Å to maintain the β-sheet structure in the I45F and T48P active conformations.

### 3. Supporting Tables

**Table S1:** List of amino acid residues constituting the S1’, S2’ and S3’ subpockets in the representative wildtype active, I45F active, T48P active and M51F shifted active conformations of γ-secretase obtained from the corresponding GaMD simulations. The residues that are within 5 Å of APP substrate residues P1’, P2’ and P3’ are listed in the table.

| System | S1’ | S2’ | S3’ |
|---|---|---|---|
| **Active (Wildtype)** | V261<br>L268<br>R269<br>L271<br>V272<br>L381<br>G382<br>P433 | V272<br>I287<br>V379<br>K380<br>L381<br>G382<br>L425<br>K430<br>A431<br>L432 | Y154<br>L271<br>V272<br>A275<br>T281<br>L282<br>F283<br>I287<br>G378<br>V379<br>K380<br>L381<br>G382 |
| **Shifted (M51F)** | I143<br>T147<br>L150<br>Y256<br>D257<br>A260<br>V261<br>L271<br>V272<br>A275<br>E276<br>P433 | D257<br>V261<br>R269<br>V272<br>E273<br>Q276<br>E277 | V261<br>V379<br>K380<br>L381<br>Y389<br>L421<br>T422<br>L425<br>A431<br>L432<br>P433<br>A434<br>L435 |
| **Active (I45F/T48P)** | V261<br>V272<br>F283<br>I287<br>K380<br>L381<br>G382<br>D385<br>A431<br>L432<br>P433<br>A434<br>L435 | V82<br>L85<br>V261<br>V379<br>K380<br>L381<br>G382<br>D385<br>L418<br>T421<br>L422<br>L425<br>A431<br>L432<br>P433<br>A434<br>L435<br>P436 | V272<br>A275<br>L282<br>I287<br>V379<br>K380<br>L381<br>L425<br>K430<br>A431<br>L432 |
